# Fast updating feedback from piriform cortex to the olfactory bulb relays multimodal reward contingency signals during rule-reversal

**DOI:** 10.1101/2023.09.12.557267

**Authors:** Diego Hernandez Trejo, Andrei Ciuparu, Pedro Garcia da Silva, Cristina M. Velasquez, Benjamin Rebouillat, Michael D. Gross, Martin B. Davis, Raul C. Muresan, Dinu F. Albeanu

**Affiliations:** Cold Spring Harbor Laboratory, Cold Spring Harbor, NY, USA; Transylvanian Institute of Neuroscience, Cluj-Napoca, Romania; current address – Champalimaud Neuroscience Program, Lisbon, Portugal; current address – University of Oxford, UK; current address –École Normale Supérieure, Paris, France; STAR-UBB Institute, Babeş-Bolyai University, Cluj-Napoca, Romania; School for Biological Sciences, Cold Spring Harbor Laboratory, Cold Spring Harbor, NY, USA

**Keywords:** rule-reversal, piriform cortex, cortical bulbar feedback, two photon calcium imaging, PCA, multilayer perceptrons, optogenetic manipulations

## Abstract

While animals readily adjust their behavior to adapt to relevant changes in the environment, the neural pathways enabling these changes remain largely unknown. Here, using multiphoton imaging, we investigated whether feedback from the piriform cortex to the olfactory bulb supports such behavioral flexibility. To this end, we engaged head-fixed mice in a multimodal rule-reversal task guided by olfactory and auditory cues. Both odor and, surprisingly, the sound cues triggered cortical bulbar feedback responses which preceded the behavioral report. Responses to the same sensory cue were strongly modulated upon changes in stimulus-reward contingency (rule reversals). The re-shaping of individual bouton responses occurred within seconds of the rule-reversal events and was correlated with changes in the behavior. Optogenetic perturbation of cortical feedback within the bulb disrupted the behavioral performance. Our results indicate that the piriform-to-olfactory bulb feedback carries reward contingency signals and is rapidly re-formatted according to changes in the behavioral context.

## Introduction

Long-range interaction between different brain regions via feedforward and feedback signals are thought to enable flexible behaviors in rapidly evolving environments. Cortical areas are highly interconnected^1–6^, suggesting that, across modalities, information about stimuli and their behavioral significance is widely shared, even in primary sensory cortical areas^7–10^. Consequently, cortical stimulus representations can be shaped by cognitive demand and contribute to the selection and generation of actions. Within this conceptual framework, cortical feedback has been proposed to support key computations. These range from extracting fine stimulus features^11,12^ to learning associations^13,14^ and generating sensorimotor predictions^13,15–23^, to modifying and executing motor actions in accordance with behaviorally relevant goals^11,12,15,16,20,24–26,26–39^. While feedback from deeper brain regions reformats visual, auditory, and somatosensory neural representations to enable the differential evaluation of same sensory inputs^11,24^, the degree to which this is the case for olfactory processing has been less scrutinized^13,16,32,39^.

Volatile compounds bind odorant receptors in the nasal epithelium and relay odor information to the olfactory bulb (OB) glomeruli, which are sorted by odorant receptor type^40–42^. Glomerular responses are normalized and de-correlated by local circuits within the olfactory bulb^43–50^, and relayed to higher brain regions by two populations of output neurons, the mitral (MCs) and tufted cells (TCs)^38,51–62^. The major projection targets of the olfactory bulb include paleo-cortical areas such as the anterior (aPCx) and posterior piriform cortex (pPCx) and the anterior olfactory nucleus (AON), in addition to the olfactory striatum (tubercle), cortical amygdala (CoA) and the lateral entorhinal cortex (lENT)^51^. Akin to other sensory modalities, recent work revealed the existence of parallel long-range feedback olfactory processing loops^16,17,35,38,45,52,57,59,63–69^. These functional streams engage specifically the mitral and the tufted cells and their preferred cortical targets (aPCx for the mitral cells, and the AON for the tufted cells, respectively) to potentially sub-serve different computations^17,70^. In this view, the TC↔AON loop mainly represents sensory features, such as odorant identity, intensity, and timing. In contrast, the MC↔PCx loop re-shapes sensory representations to enable fine discrimination of learned odorants and sensorimotor integration as a function of behavioral contingency^17^. The primary bulbar recipients of the cortical feedback are inhibitory interneurons, in particular the granule cells (GC)^15,51,71–75^. GCs integrate feedforward sensory input, and mediate lateral and recurrent inhibition by forming reciprocal synapses with the lateral dendrites of mitral and tufted cells^15,51,76–79^. The olfactory bulb dynamically represents odor inputs based on state and context^14,45,57,63–66,68,80–82^. Cortical feedback has been a prominent candidate for shaping the odor representations of the olfactory bulb outputs. Indeed, previous work showed that cortical bulbar feedback is sparse and odor specific in naïve animals, and is strengthened by learning relevant stimuli^13,16,17,32,39^. To date, however, how cortical bulbar feedback negotiates sudden changes in stimulus reward associations, and whether it contributes to flexible updating of behavioral strategies is poorly understood.

It remains unknown whether cortical bulbar feedback relays specifically olfactory signals or conveys multimodal inputs extending beyond olfaction as a function of behavioral demands. Computational models^83–87^ based on anatomical and functional data^54,88–92^ proposed that distributed connectivity between the olfactory bulb and the piriform cortex enables long-term plasticity and sparse coding of odor identity in a concentration invariant manner, regardless of temporal variations, background, or stimulus reward value^93–97^. Furthermore, classic work suggested that the piriform cortex function extends beyond sensory feature extraction, encompassing more associative spatial orientation and contextual computations^70,94,96,98–100^. Recent functional studies reported that the anterior piriform cortex representations are largely unchanged upon learning of new stimulus-reward associations^101^. In contrast, the posterior piriform activity is modulated as a function of context, and may bind together spatial information and actions related to olfactory behaviors^102,103^.

Here we investigated whether the piriform-to-olfactory bulb feedback represents changes in the reward contingency^13,57,64,65^, in addition to odor identity information as previously reported^16,17,32,39^. Specifically, we used multiphoton imaging of calcium signals to analyze the activity of cortical bulbar feedback in expert mice performing a rule-reversal task guided by olfactory and sound cues. Our results indicate that the piriform cortex-to-bulb feedback carries multimodal reward contingency signals, and is rapidly re-formatted according to changes in the behavioral contingencies.

## Results

### A rule-reversal Go/No-Go task to assess the role of cortical bulbar feedback in behavioral flexibility

To determine whether cortical bulbar feedback supports behavioral flexibility^104,105^, we engaged mice in a rule-reversal Go/No-Go task, while simultaneously monitoring the dynamics of feedback axons and synaptic boutons via multiphoton imaging of GCaMP5/7b signals (**Figs. 1a, b**; **Supplementary** Figs. 1-3, Methods). To investigate whether feedback axons represent stimulus contingency and/or trial outcome independent of the sensory modality, we used one olfactory and one auditory cue instead of two odors. We trained water-deprived head-fixed mice to discriminate between two brief (350 ms) sensory cues: a pure tone target (‘Go’) stimulus and a monomolecular odorant (‘No-Go’). We encouraged mice to respond to the ‘Go’ stimulus by licking to collect small water rewards from a spout placed in front of their mouth. Conversely, we trained them to refrain from licking in response to the ‘No-Go’ stimulus by imposing a time-out period and delivering white-noise sound before initiating the next trial in the event of spurious licking (Methods). To disambiguate the neural signatures of reward contingency from motion artifacts related to licking and other signals, in a subset of experiments we imposed a 500 ms *delay* between the cue offset and the reporting period, when water reward was available (**Fig. 1b**, **Supplementary** Fig. 1a ‘no-delay’ vs. ‘delay’ versions). Once mice learned the stimulus-reward association and reached higher than 75% accuracy (behavioral performance), we switched the stimulus reward contingency in blocks of contiguous trials between *Odor-Go* blocks (odorant rewarded) and *Sound-Go* blocks (tone rewarded, **Fig. 1a**). No explicit cue marked the block transition (rule-reversal) events.

**Figure 1.**
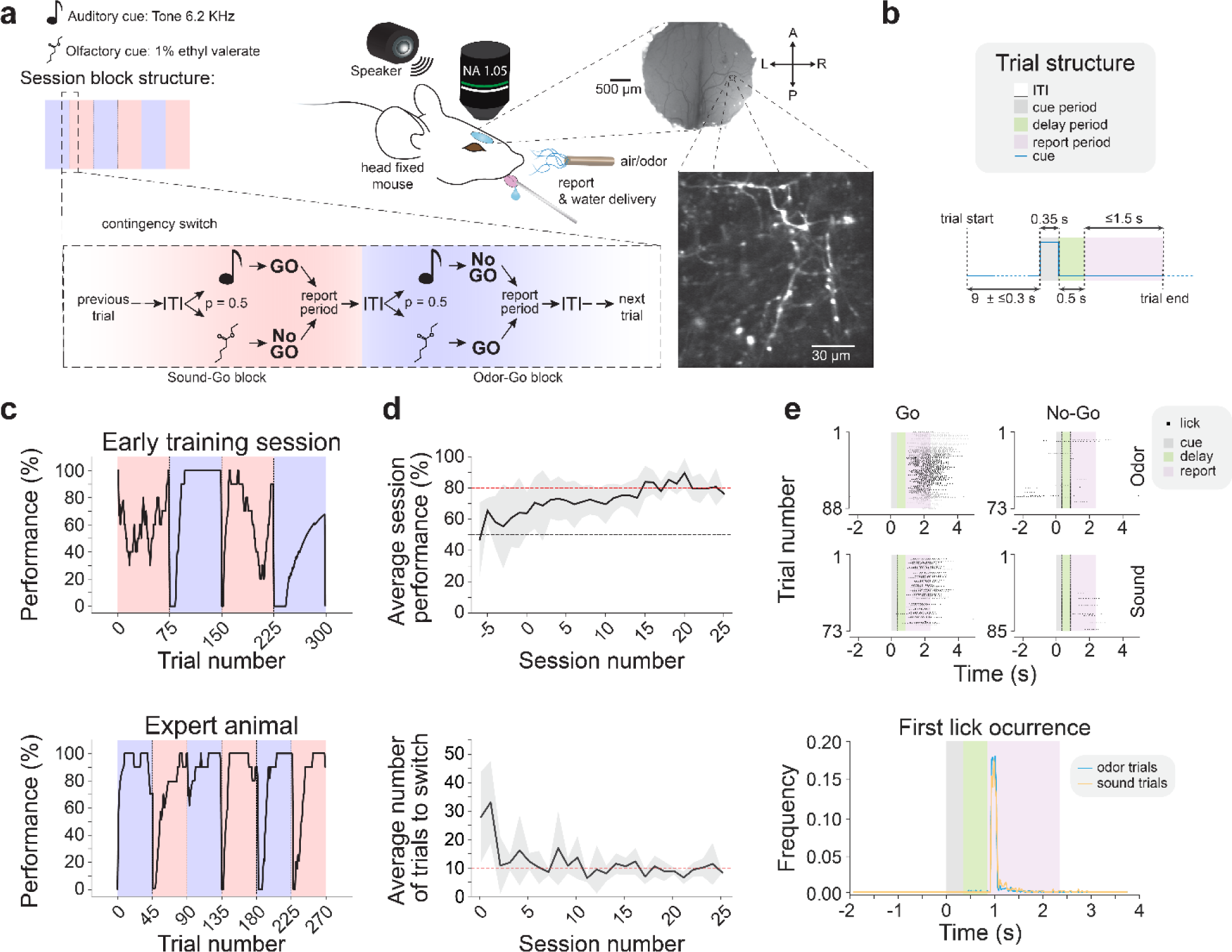
A Go/No-Go rule-reversal task using olfactory and auditory cues. (**a**) Schematics of a behavior session and example field of view. (Left) An olfactory (1% ethyl valerate) or auditory (6.2 KHz tone) cue was delivered randomly in each trial, and each session was divided into stimulus-reward contingency blocks of ∼45 trials. Stimulus-reward contingency was alternated between ‘Sound Go blocks’ (containing Sound Go and Odor No-Go trials) and ‘Odor Go blocks’ (containing Odor Go and Sound No-Go trials). (Right) Mice virally expressing GCaMP5 in the anterior piriform cortex (aPCx) with a chronic cranial window implanted above the olfactory bulb (Methods, scale bar: 500 μm). (Inset) Example field of view (FOV) of cortical bulbar feedback boutons in the granule cell layer (∼300 μm from the bulb surface, scale bar: 30 μm). (**b**) In the ‘delay’ task, a variable baseline period of (∼9 ± ≤0.3 s) was followed by the delivery of a brief odor or sound cue (0.35 s) and a fixed 0.5 s interval (delay period) before the time when the reward became available. Mice were trained to report their decision (lick vs. no-lick) within a 1.5 s window from the end of the delay period. (**c**) Example in-session behavioral performance comparisons between early training (Top) and expert (Bottom) sessions. Performance was quantified using a moving average window (bin size = 10 trials, Methods). (**d**) Progression in the behavioral performance across sessions in the delay version of the task (n = 9 mice). (Top) Average behavior session performance. Zero marks the first session when mice experienced rule-reversal in the stimulus-reward contingency. (Bottom) Average number of trials to reach 70% performance after each rule-reversal event (Methods). (**e**) (Top) Example licks (dots) from odor and sound trials (Top vs. Bottom rows) parsed by trial instruction (Go: Left; No-Go trials: Right) from one delay session. (Bottom) Distributions of latency to the first-lick from cue onset from all delay sessions parsed by cue (blue: odor; yellow: sound trials). All panels error bars: ±SEM.

Early in training, mice displayed unstable performance across blocks and were slow in updating their lick-reporting strategy upon rule-reversal (**Fig. 1c** *top*). As the training progressed (∼20 sessions), animals learned to switch reliably between reward contingencies and maintained a high level performance across the session (> 80%, **Figs. 1c** *bottom*, **d**), with drops in performance occurring only immediately after rule-reversals. Expert mice switched reporting strategies across blocks within ≤ 7 trials (6.74 ± 0.82, **Fig. 1d**, **Supplementary** Figs. 1c,d) and completed an average of 5.8 ± 0.87 reversals (blocks of ∼ 45 trials) per session, akin to other task-switching paradigms^106^. We used a stable 80% session performance as the criterion for ‘expert’ behavior and the starting point to monitor the cortical bulbar feedback activity (N = 7 mice, Methods, **Figs. 1c**-**e**). We observed comparable reaction times across modalities (930.8 ± 3.7 ms for the tone and 986.4 ± 7.6 ms for the odorant from *cue* onset in the ‘delay’ version, N= 3 mice; 445.0 ± 2.4 vs. 443.0 ± 3.2 in the ‘no-delay’ version; N= 4 mice, **Fig. 1e**, **Supplementary** Fig. 1b, Methods). The learned association was robust across days and the same animal could learn multiple odor/sound pair associations (N = 2 mice, >80% accuracy, **Supplementary** Fig. 1e). Thus, head-fixed mice mastered a rule-reversal task which enabled further investigating whether the cortical bulbar feedback supports behavioral flexibility.

### Diverse cortical bulbar feedback representations update within seconds following reward-rule switching in task-engaged mice

To monitor the cortical bulbar feedback activity in task-engaged expert mice, we expressed a genetically encoded calcium indicator in the anterior part of the piriform cortex (aPCx, EF1-FLEX-GCaMP5-AAV + AAV-Cre, Methods). We imaged fluorescence changes in synaptic boutons from feedback axons within the olfactory bulb through chronically implanted cranial windows (Methods). To determine whether this activity relates to the animals’ performance in the task, we investigated the dynamics of cortical feedback specifically during the *cue* and *delay* periods (i.e. *before licking*, Methods, N = 2,475 boutons, 20 FOVs, 3 mice, delay version; N= 1,315 boutons, 23 FOVs, 4 mice, no-delay version). Across sessions, we probed the cortical feedback activity at different depths from the surface (200-300 µm), sampling boutons mostly in the granule cell layer (**Fig. 1a**; Methods). In contrast to the sparse odor-triggered activity in naïve animals, and consistent with previous reports, we observed dense cortical feedback odor responses in the task-engaged mice^13,16^ (54.4 ± 5.0 % of sampled boutons were responsive, N = 20 FOVs, 3 mice, *delay* version; 38.3 ± 2.9 % of boutons, N = 23 FOVs, 4 mice, *no-delay* version vs. 4.3 ± 0.4 % of boutons using a 20 odors panel, N=18 FOVs, 4 naïve mice^16^, **Supplementary** Fig. 3e).

Responses typically started during the *cue* period and preceded the behavioral report (i.e. licking) (**Fig. 2a**; **Supplementary** Figs. 3-5). Surprisingly, in addition to the odor-triggered responses, changes in bouton fluorescence also occurred following the onset of auditory cues (**Fig. 2a**; **Supplementary** Figs. 3-5), revealing the presence of multimodal signals in the cortical bulbar feedback. These events were less frequent and weaker in comparison to the odor triggered activity. On average, across the cue-modulated feedback boutons, 60.0 ± 5.2 % were selectively tuned to the odor cue, 10.1 ± 2.7 % to the tone, and 29.8 ± 3.8 % responded to both types of stimuli (**Supplementary** Fig. 4a; 64.2 ± 4.7 %, 14.2 ± 3.0 %, 21.5 ± 3.2 % for the *no-delay* version, **Supplementary** Fig. 5b). To determine whether different groups of bouton responses relate to specific events throughout the trial, we performed K-means clustering of the bouton responses, independently for the odor and sound trials (Methods). Several distinct groups of feedback responses emerged which showed characteristic differences in polarity (enhanced, 47.4 ± 7.7 %, suppressed, 42.6 ± 7.9 %, complex, 9.9 ± 2.4 %; **Supplementary** Figs. 6a,c)^16,107–110^, shape, and kinetics. Some responses span mostly the duration of the cue, others slowly ramped post-cue onset and/or lingered for seconds after cue offset (**Figs. 2a-c**, **Supplementary** Figs. 4b,c, **5a**,**c**,**d**). Indeed, the time course of the responses aligned to different trial events, including the sensory *cue* (*odor* response clusters #1-3; *sound* #1, 4, 7, 8), *delay* (*odor* clusters #4, 6; *sound* #3), and *reporting* (*odor* clusters #6, 9, 10; *sound* #2, 5) periods (**Supplementary** Figs. 4b,c, **5c**,**d**). The observed diversity in feedback activity assessed via GCaMP imaging was absent in EGFP expressing boutons under same conditions in control mice (**Supplementary** Figs. 3b,d,**e)**; only a small fraction of EGFP boutons passed our responsiveness criterion, Methods (15.5 ± 7.5%; 3 FOVs; 2 mice, delay version; 0%, 4 FOVs, 1 mouse, no delay version). In addition, the number of responsive boutons and strength of response did not correlate with the frequency of licks in either task version (**Supplementary** Fig. 1f-i; GCaMP and EGFP controls). Thus, the cortical bulbar feedback activity relates to various trial events, and cannot be simply explained by motion artifacts.

**Figure 2:**
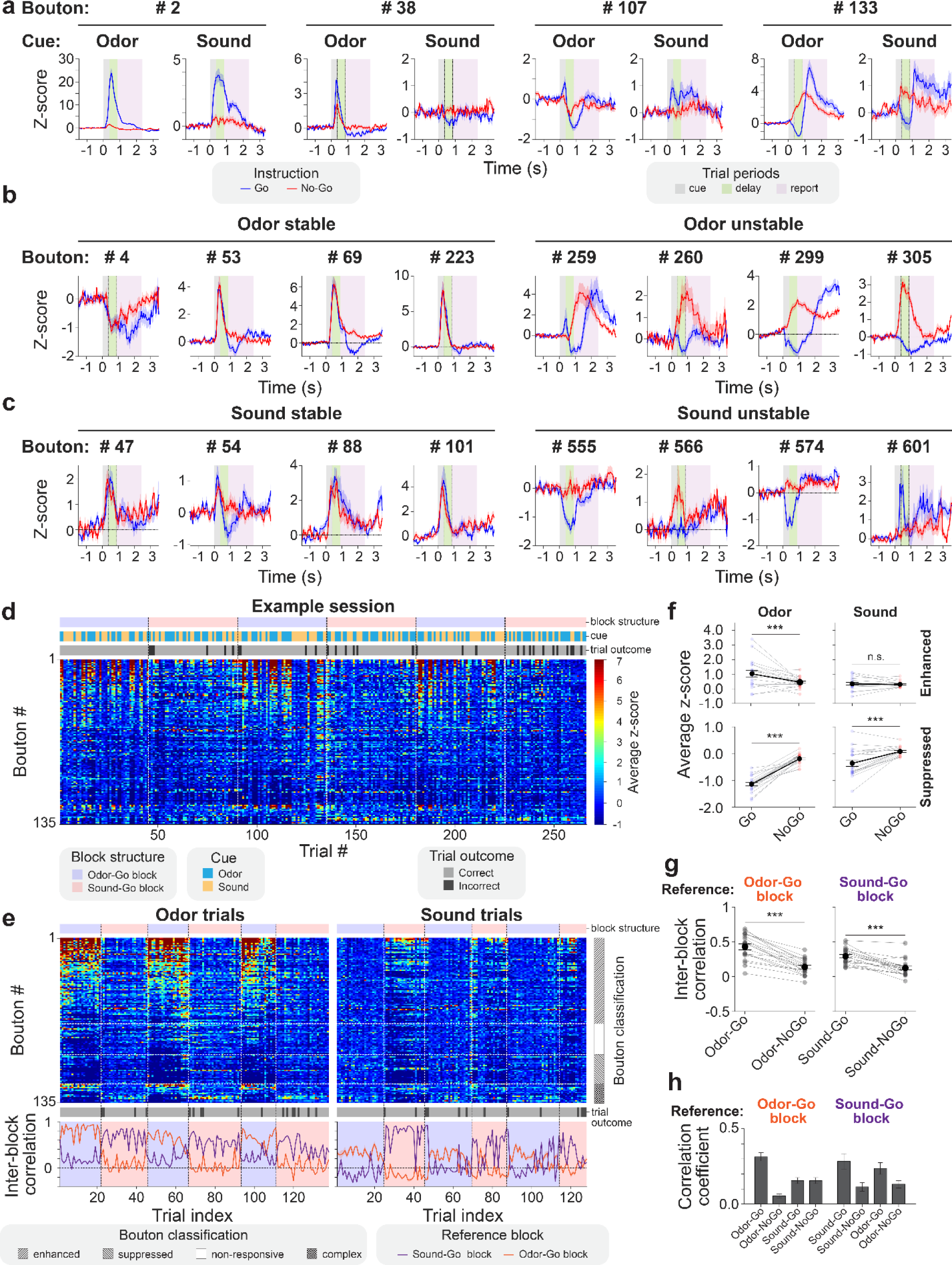
Fast update of cortical bulbar feedback representations following reward-rule switching in task-engaged mice. (**a**) Example average responses of 4 cortical bulbar boutons to odor and sound cues during Go (blue) and No-Go (red) trials. Shaded areas mark different task periods: cue (gray); delay (green); report (pink). Blue/red traces represent the average change in fluorescence (z scored) across trials; shaded area corresponds to SEM. Note the different response amplitude scales. (**b**,**c**) Example boutons that displayed stable (Left) or unstable (Right) average responses to odor (**b**) and sound (**c**) across conditions (Go vs. No-Go) assessed throughout the cue and delay periods. (**d**) Bouton responses (z-scored) averaged throughout the delay period and shown across trials in an example field of view from an expert mouse performing the rule-reversal task. Each row shows the response of one bouton across and blocks (Odor Go; Sound Go, Odor Go, etc.) throughout the behavior session. Boutons are sorted from top to bottom by the strength of their response during the odor trials. Color scale bar: Average z-score. Color-coded bars on top mark the block structure (Odor Go block vs. Sound Go block), cue identity (Odor vs. Sound) and trial outcome (Correct vs. Incorrect). (**e**) (Top). Same session as (**d**) re-sorted by cue identity: odor trials (Left) and sound trials (Right). Boutons were classified as enhanced, unresponsive, suppressed or complex (enhanced + suppressed) as per their response strength and polarity to the odor cue; (Right) same ordering of boutons was kept for the sound trials. (Bottom) Inter-block correlation analysis (Odor Go vs. Sound Go blocks): Pearson correlation coefficient calculated between the average feedback bouton ensemble response vector in the first Sound Go (purple) or first Odor Go (orange) block in the session and the ensemble bouton response vector of each trial; panels are sorted by the cue identity (Odor – Left; Sound - Right). (**f**) Average z-scored response values during the delay period parsed by cue (Odor or Sound) and instruction (Go or No-Go) for the responsive boutons sampled. Each pair of connected colored dots represents average z-scored response values computed across conditions (Go vs. No-Go) using the boutons from one behavior session. Black dots represent average z-scored ensemble bouton response across sessions. (**g**) Average inter-block correlation coefficients obtained as described in **d** for all fields of view (delay version); the first ‘Odor-Go block’ (Left) or ‘Sound-Go block’ (Right) in a session was used as reference for calculating the average feedback bouton ensemble response vector for odor and sound trials. Student’s t-test: *** = p < 0.0001. Each pair of gray connected dots represents the inter-block correlation values computed for one behavior session. Black dots represent the average correlation. (**h**) Same analysis as in (**g**) including comparisons across modalities (e.g. Odor Go trials to Sound Go trials, etc.). All panels error bars: ± SEM.

The feedback responses to both olfactory and auditory cues and their apparent alignment to different trial epochs raise the possibility that responses change flexibly, depending on the reward-rule and/or trial outcome. To test these hypotheses, we compared bouton responses to the same sensory cue across different stimulus-reward contingencies. We observed diverse responses, ranging from stimulus-tuned (odor vs. sound responsive irrespective of reward contingency; e.g. bouton #38, **Fig. 2a**), to *instruction*-tuned (‘Go’ vs. ‘No-Go’) across sensory modalities (e.g. boutons #2, #133, **Fig. 2a**). Responses of individual boutons to the same sensory stimulus often varied in shape, kinetics, and amplitude, depending on the *instruction* signal across blocks within the same behavior session (**Figs. 2b,c**; *unstable*). In contrast, other boutons were not altered by changes in reward contingency (**Figs. 2b,c**; *stable*; **Supplementary** Figs. 6e-g).

To further investigate differences in the cortical bulbar feedback activity across stimulus-reward contingencies, we used correlation analysis of bouton *ensemble* responses, as well as of *individual* bouton responses. We compared the dynamics of the feedback responses to the same sensory cue across blocks of different reward contingencies. In example field of views (**Figs. 2d,e**; **Supplementary** Figs. 7a-c), and generally across the data (**Fig. 2f; Supplementary** Fig. 2d), the odor responses appeared stronger in amplitude, and more boutons were responsive in the blocks of trials when the odor was rewarded than when it was not. In the same session, a subset of feedback boutons responded to the tone, specifically in the sound-rewarded blocks. In comparison, in blocks in which the tone was not rewarded, sound responses were less frequent and generally smaller in amplitude (**Fig. 2d,e**; **Supplementary** Figs. 7a,b; also across the data, especially suppressed responses, **Fig. 2f**). Complementary to these flexible cortical feedback representations, we also observed many boutons that responded stably to a given cue across conditions (**Figs. 2b,c**, **Supplementary** Fig. 6e-g). These boutons may enable decoding of stimulus identity, independent of its contingency. In addition, we found in control experiments that the instruction signals (Go vs. No-Go trials) modulate the sniff rate irregularly and only mildly. This modulation varied across mice and cue types (odor vs. sound) and was not correlated with the behavioral performance (**Supplementary** Fig. 8), suggesting that the reward-contingency dependence of cortical bulbar feedback activity cannot be simply explained by changes in stimulus sampling behavior.

Many boutons mirrored closely the block-structure of the task, and changed their response (shape and/or amplitude) to the same cue within seconds of each rule-reversal event (for illustration, in **Fig. 2e** we re-sorted the trials from the session shown in **2d** by Odor and Sound cues respectively). We parsed and averaged the activity of individual boutons during the *cue* (**Supplementary** Fig. 7a), *delay* (**Fig. 2d,e**), or *reporting* (**Supplementary** Fig. 7b) periods. In each of these intervals, the correlation analysis indicated that the ensemble feedback bouton responses are more similar across trials of the same block type and different across blocks of opposite reward contingency (in both versions of the task, **Figs. 2e** *Bottom*,**g**,**h**, **Supplementary** Figs. 5e *Bottom*,**f**; Methods). Similar results were obtained using self-organizing map analysis (**Supplementary** Fig. 9; Methods). Overall, within a given field of view, the ensemble cortical bulbar feedback responses appeared more similar in Go trials across modalities (Odor Go vs. Sound Go) than when compared to No-Go trials of same modality (Odor Go vs. No-Go; Sound Go vs. No-Go, **Fig. 2h**).

We further analyzed whether the time-varying fluorescence signals of *individual* boutons before reporting, throughout the *cue* and *delay* periods (during the *cue* period for the no-delay version) changed as a function of trial behavioral contingency. To this end, we compared the activity of individual boutons across hit (H) vs. correct rejection (CR) vs. false alarm (FA) trials (**Fig. 3a**). Miss (M) trials occurred rarely (< 3 %) and were thus excluded from the analysis. Overall, responses of a given bouton were similar within the same condition and diverged across different conditions (**Fig. 3a**). Boutons that differentially modulated their responses across conditions (beyond 90% percentile of the distribution of trial-to-trial variations for within condition comparisons, Methods) represented a significant fraction of the responsive population in both versions of the task (44.8 ± 10.6 % H vs. CR, 35.0 ± 7.5 % H vs. FA; 46.1 ± 10.6 % CR vs. FA Odor trials; 44.5 ± 13.1 % H vs. CR, 27.7 ± 8.4 % H vs. FA; 50.0 ± 13.5 % CR vs. FA Sound trials, **Fig. 3a**; similar for the *no delay* version, data not shown). Across fields of view, cortical feedback response amplitude during the hit and false alarm trials was generally higher than for correct rejection trials (**Supplementary** Figs. 7e,f). However, we also observed differences in the responses of individual boutons when performing pairwise comparisons between trials of different contingencies in which mice licked the reward port (**Fig. 3b**; Odor hits vs. false alarms, 65.0 ± 7.5 %; Odor vs. Sound hits, 60.1 ± 11.9 %, Odor vs. Sound false alarms, 42.7 ± 12.1%). Since in all these cases, mice subsequently licked the reward port, changes in the response of individual boutons to same stimulus across contingencies cannot be solely attributed to motion artifacts and/or preparatory motor activity. Overall, in task-engaged mice, the cortical feedback activity updated fast within the same session, depending on changes in the reward contingency.

**Figure 3.**
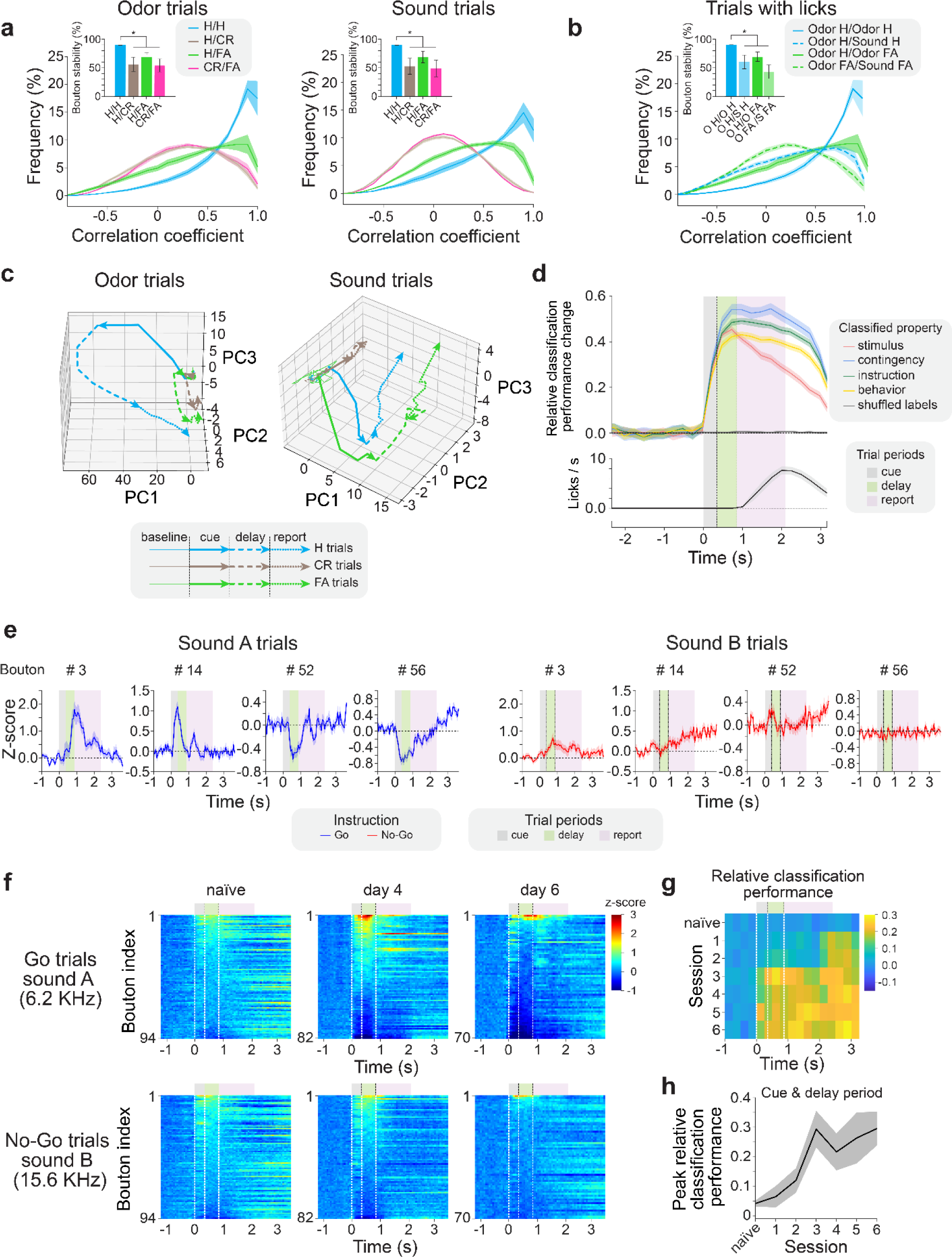
Cortical bulbar feedback represents stimulus identity, contingency, and behavioral outcome. (**a**) Histogram of individual bouton response correlation values across trials as a function of trial behavioral contingency (Hits, H vs. false alarms, FA vs. correct rejections, CR) in Odor (Left) and Sound (Right) trials. Bouton responses were sampled between cue onset and the end of delay period (before licking, Methods). Shaded areas correspond to SEM; **Inset:** Bouton response stability across conditions (H/H, H/CR, H/FA, CR/FA) reported using as reference the 90^th^ percentile of the Hit/Hit bouton response correlation distribution (bootstrap analysis, Methods). (**b**) Individual bouton response stability analysis for trials where mice subsequently licked the reward spout (hits and false alarms). Note the differences in trial-to-trial response correlation distributions when comparing Odor H/H vs. Odor H/FA vs. Odor FA/Sound FA trials. (**c**) Principal component analysis (PCA) for one example session: cortical bulbar feedback bouton ensemble response trajectories plotted in a space defined by the first three principal components (74.5 and 73.7 % variance explained respectively for odor and sound trials); population response trajectories rapidly diverge as a function of trial contingency (hits - blue vs. correct rejections - red vs. false alarms - green) for both Odor (Left) and Sound (Right) trials. Miss (M) trials were excluded from the analysis due to their low frequency (**<** 3%). Different task periods in each trajectory are represented by distinct traces (baseline: thin continuous; cue: thick continuous; delay: thick interrupted; report: thick dotted line). (**d**) Multi-layer perceptron classifiers were trained to decode stimulus identity (odor or sound), behavioral contingency (H, FA, CR), trial instruction (Go or No-Go), and behavior (lick or no-lick) in the delay version of the task. **Top:** Average classifier performance across all sessions normalized relative to baseline performance. When shuffling trial labels on the training data, the average classifier performance was 0. **Bottom:** Distribution of the number of licks per second across all sessions. (**e**) Example average responses of individual cortical bulbar bouton responses to Go (Sound A, Left) and No-Go (Sound B, Right) cues in a sound vs. sound Go/No-Go task. Shaded areas mark different task periods: cue (gray); delay (green); report (pink). Blue/red traces represent the average change in fluorescence across trials (z-scored); shaded area corresponds to SEM. (**f**) Average cortical feedback bouton responses in example fields of view parsed by instruction (Go or No-Go) and across days of training (naïve, day 4, and day 6). (**g**) Average multi-layer perceptron performance for decoding the instruction signals (Go vs. No Go) across training sessions (N=20 FOVs, 3 mice) for the task in **e**,**f**. (**h**) Peak performance of the classifier sampled from cue onset to the end of the delay period in the two sounds Go/No-Go task. All panels error bars: ±SEM.

To visualize potential differences in the feedback response trajectories as a function of behavioral contingency (H vs. CR vs. FA trials), we used principal component analysis (PCA) in individual fields of view. For systematic quantification, we further used cross-validated decoding approaches. In many arbitrarily chosen fields of view, the population trajectories for the odor, as well as the sound trials (shown in a space defined by the first three principal components, **Fig. 3c**) diverged early in the trial, typically during the *cue* period. We further trained and cross-validated classifiers (multi-layer perceptrons, MLP, Methods, **Fig. 3d, Supplementary** Figs. 10a-f, **11a-f**) to decode different task features including stimulus identity (odor vs. sound) and instruction (Go/No-Go), behavioral outcome (lick/no lick), and trial behavioral contingency (hits, H, correct rejections, CR, false alarms, FA). Of note, given our task design, many of these features are interrelated and, thus, cannot be fully assessed separately. Both in arbitrary example fields of view (**Supplementary** Fig. 10a) and when averaging classification performance across FOVs (Methods), the classifiers’ performance for decoding each of these variables rapidly increased during the *cue* and peaked during the *delay* period. The performance remained high throughout lick-reporting, as well as for several seconds after water collection (**Fig. 3d**). In the *no-delay* version of the task, decoding performance returned to baseline more rapidly than in the *delay* version, reflecting faster offset kinetics in the cortical bulbar feedback (**Fig. 3d**, **Supplementary** Figs. 11a,b).). Performance of decoders trained to discriminate between cues (odor vs. sound) decayed faster relative to the ability to report other features (Go/No-Go instructions, stimulus contingency, etc.), consistent with the transient nature of the sensory input in comparison to other trial variables analyzed (**Fig. 3d**, **Supplementary** Fig. 10a). The representation of stimulus identity appears to occur at the level of specific ensembles of cortical-bulbar neurons. Shuffling bouton labels resulted in substantial decrease in the performance of the classifiers (**Supplementary** Figs. 10c, 11e).

As expected, the classifier performance did not rise above baseline in shuffled controls, and was substantially higher in GCaMP expressing mice compared to EGFP control experiments (**Fig. 3d**; **Supplementary** Fig. 10e, Methods). Our results indicate that cortical bulbar feedback carries stimulus identity, contingency, and behavioral outcome signals, which are readily reformatted in different behavioral contexts.

Does the emergence of sound-driven cortical bulbar feedback activity require that the odor and sound cues occur in close temporal proximity? Or does it rather reflect changes in the contingency of behaviorally relevant stimuli across sensory modalities? To start answering these questions, we monitored the activity of cortical bulbar feedback boutons expressing GCaMP7b^111^ in mice trained in an auditory-only Go/No-Go task (Sound A vs. Sound B, *no odors*, Methods). During training, care was taken so that no odor-cues were present. Similar to the previous analysis, we analyzed the changes in the cortical feedback bouton fluorescence from the *cue* onset to the end of the *delay* period (prior to licking). We observed significant sound-triggered responses in naïve mice (first session on the training and imaging rig, Methods), as well as during learning, and in expert animals (**Fig. 3e**, **Supplementary** Fig. 12). Across fields of view, over the course of six days analyzed (N= 3 mice per day), the amplitude and dynamics of the sound-triggered responses changed substantially, revealing unexpected complexity (Naïve: 4.6 ± 4.6%; Day 4: 16.4 ± 2.4%; Day 6: 8.3 ± 7.7% responsive boutons). As learning of the sound-reward associations progressed, the responses of cortical feedback boutons became more tuned to the rewarded (Go) sound cue, and displayed both enhancement and suppression with respect to baseline (**Fig. 3f**, day 6; **Supplementary** Fig. 12b). Across sessions, the performance of classifiers for decoding the *instruction* signals (Go vs. No-Go) improved with training, and did not plateau within the six-days training window (**Figs. 3g-h**). As training progressed, signal *instructions* could be decoded progressively earlier within the span of a trial (during the *cue* and *delay* periods, **Fig. 3g**). Overall, these observations are consistent with the hypothesis that the cortical bulbar feedback represents, in addition to odor-specific information, reward contingency signals across different modalities.

As in our task rule-reversals occur in the absence of an overt cue, at the boundary between blocks, expert mice usually take a few trials (≤ 7, **Supplementary** Figs. 1g,h) to compute that reward contingencies have flipped and to update their reward collection strategy accordingly. This raises the question of whether cortical bulbar feedback activity mirrors the *perceived* current reward-rule, and thus lags in updating in a similar manner as the animal’s behavior performance. Indeed, we found signatures of such representational leakage in the cortical feedback bouton ensemble activity: *post* rule-reversal events, ensemble feedback activity characteristic to the previous block persisted for several trials (**Fig. 4a**). This was also reflected at the level of individual bouton responses across trials (**Fig. 4b**).

**Figure 4.**
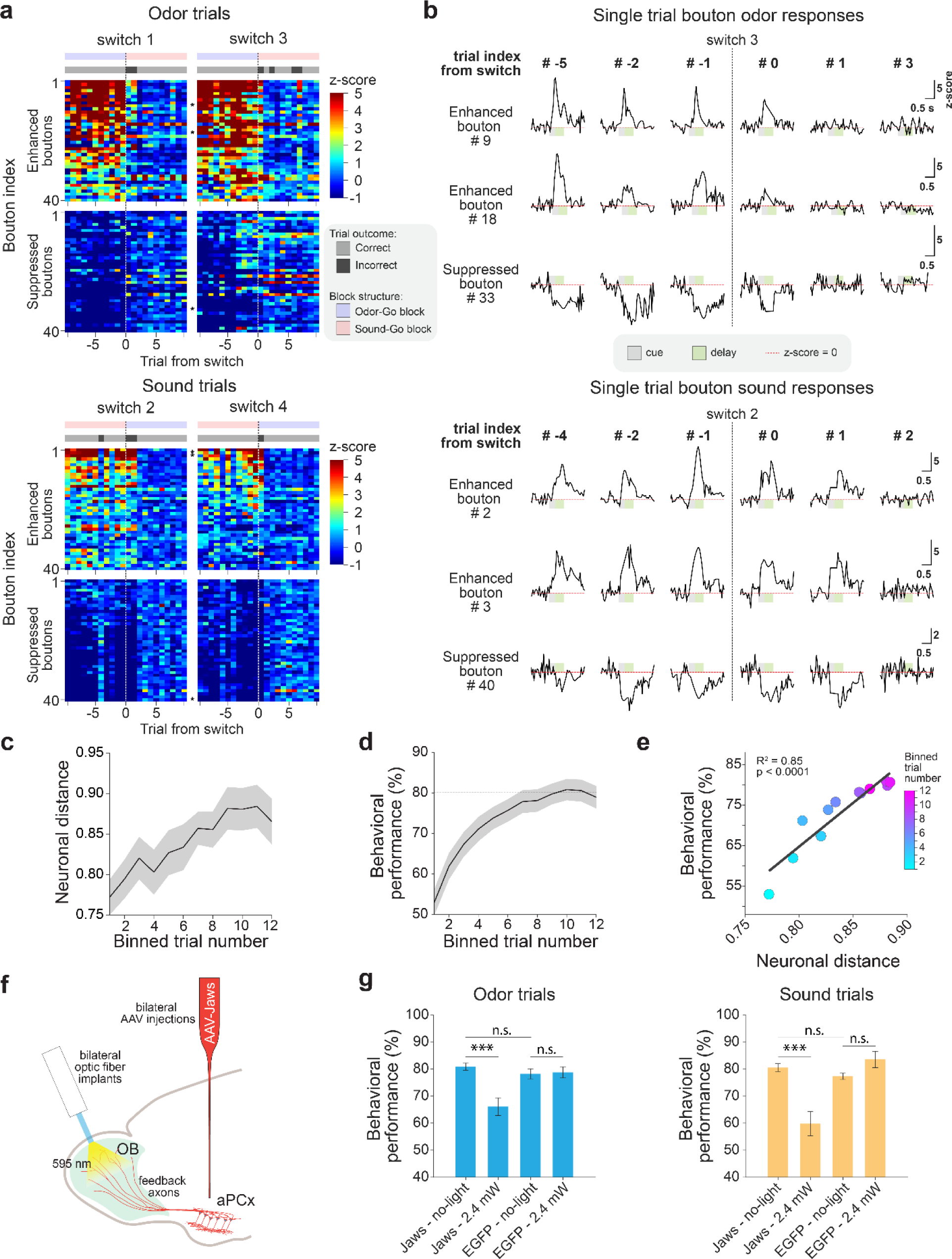
Cortical bulbar feedback activity mirrors the perceived reward-rule: (**a**) Example bouton responses during block transitions sampled throughout the ‘delay-period’ (from cue offset and before behavioral reporting) from one field of view in an expert mouse. (Top) Odor trials. (Bottom) Sound trials. Each panel shows two block transitions (columns) across 40 responsive boutons in the field of view. Gray bars mark the trial outcome (correct – light gray; incorrect-dark gray). Asterisks mark the example bouton responses shown in (**b**). (**b**) Example individual bouton response time traces from (**a**) to odor (Top) and sound (Bottom) before (negative trial index) and after (positive trial index) the contingency switch (0; vertical segmented line). Interpolated responses are shown (Methods). (**c**) Block transition neuronal distance analysis: Pearson correlation (ρ) was calculated between the cortical feedback bouton ensemble response of a given trial of the current block and the average bouton ensemble response over the last five trials of the preceding block. The neuronal distance is defined as 1 – ρ, and shown for the first twelve trials of a given block as an average across blocks and sessions. (**d**) Average behavioral performance following block transitions across sessions quantified using a three-frames sliding window. (**e**) Correlation between the neuronal distance and the block behavioral performance showed in **c** and **d**. Color bar: Trial index of each correlation value. (**f**) Optogenetic perturbation of aPCx-originating feedback performed locally within the olfactory bulb; mice bilaterally expressing Jaws in aPCx neurons were chronically implanted with optic fiber cannulas placed on top of each bulb hemisphere and trained in the rule-reversal task. In expert mice, cortical feedback was suppressed (2.4 mW, 595nm) in 25% of the trials of a behavior session. (**g**) Behavioral performance quantified for odor (Left) and sound (Right) trials independently in Jaws-aPCx and EGFP-aPCx expressing mice respectively: (Jaws – no-light) vs. (Jaws – 2.4 mW) light-on trials. ANOVA and ‘Multiple comparisons of means’: *** = p < 0.0001; n.s.: non-significant. All panels error bars: ±SEM.

We calculated a neuronal distance (1-Pearson correlation) between the bouton ensemble response (*delay* period) trajectory of the preceding block and the ensemble trajectory of each trial of the current block to the same stimulus for both odor and sound trials (Methods). This metric increased systematically across trials, and matched the increase in behavioral performance in the new block *post rule-reversal* (**Figs. 4c,d**). Plotting the mean neural distance in the ensemble trajectory versus the average behavioral performance (across blocks and sessions) revealed robust correlation (*R^2^* = 0.85, p <0.0001) between the cortical feedback activity and the update in behavioral reporting strategy (**Fig. 4e**). Qualitatively similar results were obtained when performing the analysis for odor and sound trials independently (data not shown).

To determine whether cortical bulbar feedback is necessary for expert mice to perform our task, we applied local optogenetic perturbations on the cortical bulbar feedback axons within the olfactory bulb. Using a viral strategy, we expressed Jaws, an inhibitory opsin^112^ in piriform cortex neurons (via AAV-Jaws-EGFP injections in the anterior part of the piriform cortex, aPCx, Methods; **Fig. 4f**). We monitored changes in the behavioral performance in both odor and sound blocks in catch trials during light stimulation (25% of trials, Methods). Perturbing cortical feedback activity by local optogenetic stimulation within the olfactory bulb impaired the behavior performance compared to control trials (Odor trials – Jaws-2.4 mW: 66 ± 3 % vs. 81 ± 1 % Jaws-no light; Sound trials – Jaws-2.4 mW: 60 ± 4 % vs. 81 ± 1 %; p < 0.001, N = 3 mice; **Fig. 4g**) and to sessions using mice that expressed EGFP only in the cortical bulbar feedback axons (under same light stimulation conditions, **Fig. 4g**, Methods). These differences in behavioral performance were reflected as increases in the rate of false alarms and misses for both the odor, as well as the sound trials. Our results are consistent with a scenario in which the cortical bulbar feedback contributes to assessing the behavioral (reward) contingency stimuli across multiple sensory modalities (e.g. odor and sound cues), and relays signals to the olfactory bulb that extend beyond processing olfactory input.

## Discussion

Taking advantage of a novel Go/No-Go rule-reversal task engaging olfactory and auditory cues (**Fig. 1**), we found that the feedback from the piriform cortex to the olfactory bulb relays reward contingency information across multiple sensory modalities. The cortical bulbar feedback is reformatted upon changes in stimulus-reward contingency rules, and mirrors the behavior of expert mice across rule-reversals (**Figs. 2-4**). Furthermore, optogenetic suppression experiments (**Fig. 4**) suggest that the cortical bulbar feedback is part of a larger processing network that enables mice to adapt to sudden changes in stimulus-reward contingencies.

To investigate whether the cortical bulbar feedback represents reward contingency and supports behavioral flexibility^104,105,113^, we focused our analysis on the cue-evoked responses of feedback boutons preceding the behavioral readout (lick/no-lick assay). As in previous work^16,107–110^, we observed both enhanced and suppressed feedback responses that were roughly balanced in their frequency. The presence of enhanced and suppressed responses may increase the dynamic range of cortical action in controlling the activity of bulbar outputs. Decoding analysis suggested that both the enhanced and suppressed feedback responses participate to representing various stimulus features (e.g. identity, contingency, etc., **Supplementary** Figs. 10b, 11c). Further analysis is, however, necessary to determine whether these signals, arising presumably from distinct populations of piriform outputs^107–110^, carry signals involved in different computations^16,23^.

To increase the separability of potential motion artifacts and rule-related signals, we imposed a short delay between the offset of the sensory cue and the behavioral reporting. Many feedback boutons modulated their responses in tight correlation with changes in the reward contingency rules (**Figs. 2**,**3**). The re-organization of cortical feedback activity included re-shaping the kinetics, amplitude, and response polarity of individual boutons (**Figs. 2**, **3**; **Supplementary** Fig. 6). It generally lagged the rule-switching events by a few trials (∼7; **Supplementary** Fig. 1) and was correlated with changes in behavioral performance (**Figs. 2**, **4**). Responses in the piriform-to-bulb feedback were triggered by both the odor and the sound cues **(Figs. 2**, **3**, **Supplementary** Figs. 4, **5**). Feedback bouton activity modulation across conditions (Go vs. No-Go blocks) could not be simply explained by motion artifacts as indicated by EGFP control experiments (**Supplementary** Figs. 1f,i; **3d**,**e**;**10e**), nor by changes in sniffing (**Supplementary** Fig. 8), consistent with recent reports in related tasks^114,115^. In parallel, many bouton responses were robust to the rule-reversals, and may enable stable representations of the sensory input despite changes in reward contingency (**Fig. 2**; **Supplementary** Fig. 6). Overall, population based decoding analysis indicated that the cortical bulbar feedback carries signals related to stimulus identity, reward contingency, and trial outcome (**Fig. 3**). On average, feedback responses triggered by the same cue, had higher amplitude during hits and false alarms than during correct rejection trials (**Supplementary** Figs. 7e,f). However, we also observed differences in the responses of individual boutons across trials of different contingency in which mice subsequently licked the reward port (Odor Hit vs. Odor FA vs. Sound Hit, etc., **Fig. 3b**). Thus, the cortical feedback responses cannot solely be explained as motor preparatory activity. The decoding analyses were successful even when zooming into arbitrarily chosen individual (∼ 50 x 50 µm) fields of view, revealing robust representations of the task features in the cortical feedback activity. Across multiple rule-reversals within the same session, the neural ensembles transitioned fast back-and-forth between rule-associated representations, akin to reports in other brain regions^106,116–119^. A given stimulus triggered similar cortical feedback activity in blocks of trials of the same contingency rule, and dissimilar representations in blocks of the opposite reward contingency, revealing attractor-like behavior in the piriform-to-bulb neural dynamics. Furthermore, Go feedback ensemble responses across modalities (Odor Go vs. Sound-Go) appeared more similar than responses to the same cue across instruction signals (e.g. Odor Go vs. Odor No-Go; Sound Go vs. Sound-No Go, **Fig. 2h**).

The fast updating of the piriform cortex-to-bulb feedback responses upon rule-reversal contrasts previous reports that anterior piriform cortex representations are stable, largely sensory, and only mildly modulated by learning, context, and rule reversal^100,101^, but see^98^. Differences in the behavioral tasks employed across studies, and potential specificity in the activity of distinct piriform output cell types defined by their long-range projections, may account for these differences. Indeed, different groups of piriform output neurons target functionally distinct brain regions (olfactory bulb vs. orbitofrontal cortex vs. cortical amygdala vs. lateral entorhinal cortex, etc.), and are enriched at distinct locations along the anterior-posterior axis^120–122^. To date, however, most studies monitored activity in the anterior piriform cortex agnostic of the projection targets of the recorded neurons^90,101,110,123–126^. As such, bulb-projecting piriform cells appear to flexibly update their representations in conjunction with changes in stimulus-reward contingency. In contrast, other piriform output neurons that target, for example, the orbitofrontal cortex or other brain regions may be less affected by stimulus-reward associations, and primarily represent sensory features of stimuli^101^.

Our results open venues for investigating the mechanisms supporting the flexible gating of piriform-to-bulb feedback signals. Since the changes in response amplitude and kinetics occur within seconds (a few trials from rule-reversal), they may rely on fast gating signals, rather than slower synaptic plasticity-based changes. Further investigation is necessary to determine whether these signals originate in the piriform cortex, or emerge through interactions with other association cortical areas (e.g. OFC, mPFC, lENT)^114,127,128^, and/or reflect neuromodulatory action^129–132^ onto specific piriform circuits^133–135^. While calcium dynamics in axons terminals have been shown to reflect changes in firing rates at the soma^16,26,136^, an alternative possibility is that the gating of calcium signals in the cortical feedback axons occurs via interneuron input within the bulb. However, the presence of sound-evoked responses in the feedback boutons, modulated by reward contingency suggests that a local (bulbar) mechanism is not a parsimonious explanation.

The activity of mitral cells is modulated by context, learning, and stimulus contingency, and cortical bulbar feedback has been singled out as a potential signal responsible for shaping these bulb outputs^13,45,57,63–68,80,81,137^. A recent study reported reward-related signals in the mitral cells which are modulated by the piriform cortex-to-bulb feedback (assessed via pharmacological silencing of the piriform)^115^. Our data is consistent with this body of work; it provides a framework to further analyze the dynamics of mitral (and tufted) cells in mice engaged in rule-reversal tasks, and under more naturalistic conditions^138^, in the presence and absence of cortical feedback. Parallel feedback loops engaging the mitral vs. tufted cells and their dominant cortical targets, the piriform cortex and AON have been reported to perform different computations^17^. For example, odor identity and concentration are more easily read-out from the tufted cell ensemble representations, whereas mitral cells may represent subtler features of odorants^17,57,65,80,81^. We expect mitral cell activity to change within a few trials post-rule reversals, matching the re-organization of cortical feedback responses and changes in behavioral performance (i.e. lower mitral cell response amplitude in the Go vs. the No-Go trial blocks). Further, we expect that the tufted cell and AON-to-bulb feedback representations are more sensory in nature, robust to changes in the stimulus-reward associations, and are less affected by perturbations of the piriform-to-bulb feedback^16,17^.

We observed sparse, but diverse, enhanced as well as suppressed sound-evoked responses in the piriform-to-bulb feedback. While sound-triggered activity has been reported in the piriform cortex^139^, its origin and computational roles remain unclear. In auditory and visual processing, inputs from the somatosensory^140,141^ and auditory cortex^7^ are thought to shape cortical neural representations as a function of experience. Specifically, these signals may relay stimulus association memories for V1 cortical circuits to compare the predicted and actual sensory inputs^7^. In contrast, a recent report^142^ in awake *passive* mice identified auditory cortex-independent, stereotyped low dimensional sound-triggered signals in the visual cortex. These responses could be predicted from small body movements and may reflect changes in internal brain states. In our experiments, the sound-triggered piriform feedback responses were apparent in expert mice engaged in the rule-reversal task (**Fig. 2**), in naïve individuals, as well as in mice performing a two-tones Go-No/Go task (**Fig. 3**). Stimulus-reward associations modulated the response kinetics and amplitude of the sound-triggered responses within the same session upon rule-reversals and during learning of the stimulus-reward associations in the two tones task. While, sound responses were diverse and aligned well to different epochs of the task (*cue*, *delay*, *report*), we did not investigate here whether varying sound stimulus features (frequency, amplitude, etc.) impacts the cortical bulbar feedback activity. Many sound and odor-evoked feedback responses in the expert mice lingered for seconds even after the reporting period, raising the possibility that they serve as lasting memory traces associated with the stimulus reward contingency^16^ (**Figs. 2**, **3**; **Supplementary** Figs. 4,5,**10**,**11**).

Why might the olfactory bulb need to “know” when a stimulus (odorant and/or especially a non-olfactory stimulus, such as a tone) is rewarded? One possibility is that the cortical bulbar feedback serves as a binding motif in cases where odors and sounds are required together for obtaining reward. While it may be more intuitive for the binding to occur further downstream, other reports have shown that information regarding one sensory type is distributed across sensory and motor pathways of another type^7–9,143–145^. An alternative possibility is that the feedback helps perform credit assignment. If subtle aspects of odorants drive the reward, these features would be more readily accessible to the cortex *post* mitral cell activity re-shaping due to cortical feedback input. Thus, a two-part representation — “what is it?” (via the TC↔AON pathway) and “what is new/different/important about it?” (via the MC↔PCx pathway) — may enable better representation of what aspects of stimuli in the environment lead to rewards. This process can be viewed as a representation learning algorithm which learns a mapping from sensory inputs to a (latent) feature space; the mitral cells may pass on a residual representation (i.e. reconstruction errors^13,15,17,19,56^), which highlights, in addition to identity, aspects of stimuli related to their associated reward contingency, context and/or level of engagement. The impaired behavioral performance observed in both odor and sound trials upon optogenetic suppression of the cortical feedback locally within the bulb (**Fig. 4**) is consistent with this scenario.

Previous work indicated that the medial prefrontal, orbitofrontal cortex, and basolateral amygdala circuits support behavioral flexibility in olfactory processing^100,101,103,146,147^. Dense orbitofrontal-to-piriform cortex bidirectional interactions^127,148^, top-down inputs from mPFC to AON, piriform cortex and olfactory striatum (tubercle), and neuromodulatory signals may shape the representations of sensory stimuli as a function of learned odor-reward associations^149–151^ and attentional state^114,128^. Our results suggest that the piriform-to-bulb feedback acts as part of a larger processing network that relates stimulus-reward associations to implementing decisions that drive behavior in dynamic environments.

## Supporting information

Hernandez_Trejo_Supplemental Information_rule_reversal

## Acknowledgements

The authors acknowledge A. Banerjee, A. Koulakov, S. Navlakha, S.D. Shea, P. Villar, H. Xu and members of the Albeanu lab for critical discussions, and R. Eifert and P. Gupta for technical support. This work was supported by NSF IOS-1656830 to D.F.A and R.C.M., CIRCUITGENUS (RO-NO-2019-0504), ERA-NET-FLAG-ERA-ModelDX Consciousness, ERANET-NEURON-Unscrambly, and NEUROTWIN (GA 952096) to R.C.M., and NIH R01DC014487-03 to D.F.A.

## Data and code availability

All data matrices included in the analyses presented here representing cortical bulbar GCaMP responses triggered by odor and sound cues in task engaged mice, EGFP controls and the behavioral performance in the rule-reversal and two-tones tasks, as well as the code used for analysis are available upon request.

## Authors contributions

Investigation, methodology, formal analyses, visualization, data curation, validation, software and writing, D.H.T., A.C.; conceptualization, investigation, methodology, formal analysis, visualization, software, data curation, P.G.S.; investigation, data curation, visualization, validation, software, C.M.V., B.R., M.D.G., M.B.D; methodology, writing, supervision, project administration, and funding acquisition, R.C.M.; conceptualization, methodology, writing, supervision, project administration, and funding acquisition, D.F.A.

## Methods

### Mice

Overall, 20 B6/129 mice were used. 11 for the rule-reversal task experiments, including 4 – *no delay*, 3 – *delay* versions of the task), 4 for the EGFP controls, 3 for the sound-sound Go/No-go task; 6 for the optogenetic suppression experiments (3 for Jaws and 3 for EGFP controls). All animal procedures conformed to NIH guidelines and were approved by the Animal Care and Use Committee of Cold Spring Harbor Laboratory.

### Surgical procedures

Mice were injected with NSAID Meloxicam 0.5 mg/Kg (Metacam®, Boehringer Ingelheim. Ingelheim, Germany) 24 hours prior the surgical procedure, at the onset of surgery, and for 2 days *post* each surgical procedure. Depending on the recovery progression, the NSAID treatment was maintained until mice showed alert, and responsive behavior. Before each stereotaxic surgery, mice were anesthetized with 10% v/v Isofluorane (Cat# 029405. Covetrus. Portland, ME, US). For the chronic window and headbar implantation procedures, mice were anesthetized with a ketamine/xylazine (125 mg/Kg - 12.5 mg/Kg) cocktail. During surgery, the animal’s eyes were protected with an ophthalmic ointment (Puralube®. Dechra. Nortwich, England, UK). Temperature was maintained at 37 °C using a heating pad (FST TR-200, Fine Science Tools. Foster City, CA, USA). Respiratory rate and lack of pain reflexes were monitored throughout the procedure. Chronic window implant surgeries were supplemented with dexamethasone (4 mg/Kg) to prevent swelling, enrofloxacin (5 mg/Kg) to prevent bacterial infection, and carprofen (5 mg/Kg) to reduce inflammation.

### Viral infections

To target the piriform cortex-to-bulb feedback for the imaging and optogenetic perturbation experiments, we used the following viruses: AAV2/9-EF1a-Flex-GCaMP5 and AAC9-Cre (Penn Vector Core, Philadelphia, PA, USA), AAV1-syn-jGCaMP7b-WPRE (Cat# 104489-AAV1), AAV1-hSyn-EGFP (Cat# 504650-AAV1), and AAV5-hSyn-Jaws-KGC-GFP-ER2 (Cat # 65014-AAV5) from Addgene (Watertown, MA, USA).

### AAV stereotaxic injections

Adult mice (males and females > 60 days old, 25-40 g) were used for the stereotaxic surgery. After the induction step (isoflurane), mice were positioned in the stereotaxic device which was fitted with a isoflurane delivery mask (Cat# 942. Kopf®. Tujunga, CA, USA), and their eyes covered with ophthalmic ointment. Head surface was cleared using hair removal cream (NairTM) and cleaned with betadine and saline. Skin was removed and the surface of the skull scraped of connective tissue to identify the cranial sutures. These were further used to align the anterior-posterior head angle by leveling the bregma and lambda with respect to the horizontal. Mice were injected bilaterally with AAV (∼280 nL per site) using a calibrated borosilicate glass micropipette (tip diameter, ∼10 μm) through small craniotomies (∼1 mm) spanning 1 mm along the anterior-posterior axis of the aPCx in both hemispheres. Coordinate 1: anterior-posterior, *AP +2.5 mm,* medial-lateral, *ML ±2.2 mm,* dorsal-ventral, *DV -3.00 mm*; Coordinate 2: *AP +2.0 mm, ML ±2.2 mm, DV -3.5 mm; Coordinate 3: AP +1.5 mm, ML ±2.8 mm, DV -3.75 mm*. AP and ML coordinates were estimated from bregma, and all DV coordinates were measured from the pia surface. AAV Injections were delivered using a Picospritzer III (General Valve) and pulse generator (Agilent) by pressure application (5-20 psi, 5-20 ms at 0.5 Hz). Mice were injected with a 1:1 mixture of AAV2/9-EF1a-Flex-GCaMP5 and AAC9-Cre for imaging, with AAV5-hSyn-Jaws-KGC-GFP-ER2 for optogenetic suppression, and with AAV1-hSyn-EGFP for the EGFP control experiments. Mice trained in the two sounds Go/No-Go task were injected with AAV1-syn-jGCaMP7b-WPRE.

### Chronic implantations

After recovery from the stereotaxic surgery, mice were implanted with a custom titanium head bar attached with C&B Metabond Quick adhesive luting cement (Cat# S380. Parkell. Edgewood, NY, USA), followed by black Ortho-JetTM dental acrylic application (Cat# 1520BLK. Lang. Chicago, IL, USA) and with a cranial window on top of the olfactory bulb as previously described ^16,17^. Special care was taken to remove the bone under the inter-frontal suture and remove small bone pieces at the edges. During surgery, the exposed olfactory bulb was continuously protected and cleaned of blood excess using artificial cerebrospinal fluid (aCSF) and aCSF-soaked gelfoam. Once both hemibulbs were exposed and clean, a fresh drop of aCSF was placed on top, followed by a 3 mm round cover glass, which was gently pushed onto the OB surface to minimize motion artifacts in further experiments. Once in place, the coverslip was sealed along the edges with a combination of VitrebondTM, Crazy-GlueTM, and dental acrylic to cover the exposed skull. Mice recovered for ∼ 7 days before imaging experiments.

### Behavioral training

Mice were water-deprived until reaching 85% of their original weight. Once the desired weight was achieved, head-fixed mice were trained to discriminate between two brief (350 ms) sensory cues: a pure tone (*6.221 KHz, 70 dB*) target (Go) stimulus and a monomolecular odorant (*1% ethyl valerate*) distractor (No-Go). The pure tone choice was based on a prime number to minimize harmonics. Care was taken to choose an odorant cue that had high SNR photoionization device (PID, Aurora Scientific) readings. Olfactory cues were presented from an odor port placed in front of the mouse’s snout and auditory cues were delivered from a speaker on the side. Each training session composed of ∼ 270 consecutive trials separated by a variable inter-trial interval (ITI; 9 ± 0.3 to 1.2 s). Trials were randomly assigned (p = 0.5) as odor trials or sound trials. In the *Go* trials, mice were trained to report the presence of the *Go* stimulus by licking a spout in front of their mouth, from which they received a small water reward (3.3 μL, *Hit* trials). In addition, mice were trained to refrain from licking (*Correct Rejections*) in the No-Go trials, so they could move faster to the next trial. Incorrect trials (*False Alarms* and *Misses*) were punished by lack of reward and the addition of a time-out period (10 s) to the regular ITI plus a one second long 70 dB white-noise sound.

Once mice learned the stimulus-reward association to higher than 75% accuracy, we switched the reward contingency in blocks of contiguous trials between Odor-Go blocks (odor rewarded) and Sound-Go blocks (tone rewarded). Within the same session, mice sample multiple rule-reversals in blocks of ∼ 45 trials. The initial stages of the rule reversal training consisted of classical conditioning to the sound cue by repeatedly pairing the sound cue to a free water delivery. In this phase, animals were allowed to lick at will, without punishment. If the animals reliably licked to the cue, we introduced the odor cue as the unconditioned stimulus. We delivered water rewards when the mouse licked for the correct cue within the designated *response* period. Once mice reached > 75% correct performance, we started reinforcing the timing parameters by turning on punishment (air puff), which was delivered for both incorrect choices and early licks, but not *Miss* trials (failing to lick for a rewarded cue). Once the animals retained > 75% correct performance, we reversed the reward contingencies, and started rewarding the odor cue. Care was taken to not over train the animals before this point. The reversal was not cued, but during the initial stage of the reversal training we switched off the timeout punishment, while instead relying on a white noise error signal. The best behavioral performance was obtained in this configuration, potentially because mice used the white noise as a general error signal. During this period of early training, the error rate was high, mice predominantly licking for the sound cue, and ignoring the odor cue. We let mice learn at their own rate, allowing as many sessions as necessary to reach the same 75% performance criterion as before the reversal. Further, we introduced rule reversals within the same session on a regular basis, while shortening the number of trials in between reversals to a steady state of ∼ 45 trials between reversals (block design). Mice were trained in one or two sessions per day, with each session lasting from 45 minutes to 1.5 hours.

We trained mice in two versions of the task. In the *no-delay* version, a trial started with a variable length *baseline* (9 ± ≤ 0.3 s), followed by the delivery of a brief sensory *cue* (0.35 s) and a reporting period (1.5 s) which started at the *cue onset*. In the *delay* version of the task, an addition 500 ms *delay period* was introduced between the *cue offset* and the start of the *reporting* period. To receive water reward in the *Go* trials, mice had to lick the water spout during the reporting period. Any trial where one or more licks occurred before the start of the *reporting* period was classified as an *early-lick* trial and excluded from further analysis. Upon achieving a steady 80% average session performance, mice were used for chronic multiphoton imaging sessions and optogenetic suppression experiments.

### Monitoring sniffing

We monitored sniffing in control expert mice performing the rule-reversal task. We used a mass airflow sensor (Honeywell AWM3300V) mounted into a 3D-printed nose mask coupled to the odor delivery system^125^. Signals were further amplified, digitized (1 KHz) and low-pass filtered (10 Hz cut-off).

### Sound-sound Go/No Go task

Mice expressing GCaMP7b in the anterior piriform cortex-to-olfactory bulb feedback were trained in a two sounds Go/No-Go task (6.2 and 15.6 KHz tone cues, at 70 dB, 350 ms). Two mice were trained with the 6.2 KHz-Go / 15.6 KHz-No-Go rule and one with the opposite rule (15.6 KHz-Go / 6.2 KHz-No-Go). Naïve response sessions were acquired before water-depriving the mice for behavioral training. The trials followed the same structure as described above for the *delay* version of the rule-reversal task.

### Multiphoton imaging

A Chameleon Ultra II Ti:Saphire femtosecond pulsed laser (Coherent) was coupled to a custom built multiphoton microscope. The shortest optical path was used to steer the laser to a galvanometric mirrors (6215HB, Cambridge Technologies) based scanning system. The scanning head projected the incident laser beam (930 nm) through a scan lens (50 mm FL) and tube lens (300 mm FL) so as to fill the back aperture of the objective (Olympus 25X, 1.05 NA). A Hamamatsu modified H7422-40 photomultiplier tube was used as photo-detector and a Pockels cell (350-80 BK, 302RM driver, ConOptics) as laser power modulator. The current output of the PMT was transformed to voltage, amplified (SR570, Stanford Instruments), and digitized using a data acquisition board that also controlled the scanning system (PCI 6115, National Instruments). Image acquisition and scanning were controlled using custom-written software in LabView (National Instruments). Using submicroscopic beads (0.5 µm) and a 1.05 NA, 25X Olympus objective, the point spread function (PSF) was calculated x-y (1.0 µm FWHM) and z (2.0 µm FWHM). Cortical bulbar feedback activity was sampled at 16 Hz (160 x 128 pixels, FOV size 48 x 48 μm, 0.30-0.38 µm pixel size) for ∼ 6 seconds per trial. Sound produced by the galvo-mirrors at the scan frequencies employed fell out of the mouse audible range (< 1 KHz), and was therefore unlikely to provide extra cue for solving the task. Mice were imaged between 4-to-6 weeks after the AAV injections.

## Data analyses

### Movement correction and ROI selection

Rigid registration in MATLAB was applied to the fluorescence time-lapse stacks acquired. Images were visually inspected to select a motion-free sequence of frames and create a median reference image to which we registered each image stack corresponding to a given trial. ROI selection was performed manually (ImageJ): we used both the median and standard deviation projections of the registered images to draw ROIs around the cortical feedback axonal boutons in the FOV.

### Bouton and frame rejection, and interpolation

We iteratively searched for an optimal interpolation method that could be applied to the data, such that we could maximize the number of trials and ROIs to be included in the further analysis steps, without introducing erroneous data. Based on our testing (see below), we identified thresholds for discarding trials and specific ROIs. Multiphoton imaging in awake head-fixed mice frequently displays signal loss due to brain motion: field of view (FOV) loss, where signal in all boutons is compromised at the same time, and bouton loss, where one ROI moves out of the plane of imaging/field of view due to a more complex movement (e.g. rotation). To find an appropriate interpolation method, we used behavioral sessions that had the fewest number of skipped frames. Specifically, we used as ground truth trials that had at least 90 contiguous frames without NaNs to evaluate the performance of several interpolation methods. We introduced NaN values across sequences of images of varying length (1 to 45) at all possible time points and analyzed the performance of three standard (*linear*, *spline*^152^, *Akima*^153^), and of one custom interpolation method, we call *step* interpolation, where we replace the missing data with the value immediately preceding it. By evaluating the mean squared error between the ground truth signal and the interpolated version, we identified the best interpolation method among the ones analyzed as a function of the number of missing frames: *Akima* (if the window of missing frames is shorter than 7), *step interpolation* (if the fluorescence value occurring just before the missing window is smaller than the value following immediately after, and the difference is greater than a set threshold); and *linear interpolation* in all other cases. For our data set, this algorithm could not reliably interpolate windows of missing data longer than 15 contiguous frames (error exceeded 1 standard deviation of the signal at windows longer than 20 frames). Using a conservative threshold, if any trial had a loss window in any of its ROIs longer than 15 frames, it was not considered for further analysis. In addition, we set a threshold of 70% for the minimum amount of valid data points in a trial: if more than 30% frames were NaNs, the trial was discarded). After trial and ROI rejection, the interpolation strategy described was applied to fill in any missing data.

### Bleaching correction

Post individual ROI extraction, we applied a bleaching correction, assuming that: (1) bleaching follows a similar trend across trials for each individual ROI; (2) during the baseline period, the measured activity is random. Given these assumptions, by averaging across trials the fluorescence of an ROI, the only robust trend potentially present in the baseline would be the characteristic bleaching for that ROI. We checked that the baseline fluorescence of a given bouton is comparable across trials within a session. The procedure used for each ROI was as follows: (1) we averaged the fluorescence traces across all trials for that ROI (aligned to stimulus presentation), (2) fitted an exponential decay function to the baseline period of the fluorescence trace; (3) extended the fitted function to the length of each trial, (4) subtracted the fitted function from the ROI trace.

### Normalization

For the decoding analysis, we normalized the data so as to use the same network across all datasets and ROIs, since multi-layer perceptrons (MLP) are sensitive to the range of the input values. For each ROI and trial, we z-scored the traces individually, using the signal baseline period to compute the necessary statistics (mean and standard deviation).

### Bouton responsiveness and classification

To evaluate the responsiveness of each bouton, we obtained the distribution of average z-score values from the baseline periods and used a 99 percentile value of the distribution as criterion to decide if a bouton was responsive. The baseline reference distribution was obtained from average z-score values each quantified over six-frames intervals extracted from the baseline period and accumulated across all trials. For each bouton and trial type, we also obtained average z-score values of equivalent six-frames periods extracted between the start of the *cue* delivery to the end of the *response* period (parsed by *cue*, *delay* and *response* periods). For each bouton and trial type (Go/No-Go), we compared the average response across trials (parsed by cue, delay and response periods) with the baseline reference distribution. If at least one of these values crossed the 99^th^ percentile of the baseline distribution criterion, the bouton was classified as responsive (*enhanced* or *suppressed*). If some of these values were under and some above the 99th percentile of the baseline distribution, the response was classified as *complex*.

### Clustering analysis

Waveforms were normalized by the largest magnitude response of the absolute value of each trace. We used the k-means clustering function in MATLAB (Euclidean distance). Cluster quality was assessed by calculating the average distance between the average waveforms assigned to each cluster (*d*). To determine the total number of clusters, we calculated the average *d*, while varying the number of clusters from 2 to 100. The total cluster number was chosen using a cutoff threshold for which the average decrease in *d* plateaued.

### Inter-block correlation analysis

To determine whether bouton response patterns change accordingly to each rule reversal, for each field of view, we used the average z-scored fluorescence values aggregated across the *delay* period for both the odor and sound cue trials (**Fig. 2e**). We assembled *an average response vector* of the 1st or 2nd block of trials within a session for each cue by randomly picking half of the trials of each of those blocks (*reference vectors*). We further calculated the Pearson correlation values between these reference vectors and each column (trial) of the *delay period response matrix* for the odor and sound trials respectively. For each session, we repeated the bootstrapping of random trials to generate reference vectors one hundred times. One example (mean ± SEM) correlation trace is shown at the bottom of the **Fig. 2e** panel. We further calculated an *inter-block correlation* by obtaining the mean ± SEM of the Pearson coefficient values of each Odor-Go, Odor-No-Go, Sound-Go, and Sound No-Go blocks for each session and mouse (**Figs. 2g,h; Supplementary** Fig. 5f).

### Bouton stability

To evaluate how stable individual bouton responses were across trials of different stimulus-reward contingencies, we calculated the distributions of pairwise Pearson correlation values within (e.g. Hits-Hits) and across different (e.g. Hits-False Alarms; Hits-Correct Rejections, etc.) reward contingency conditions (**Figs. 3a,b**). Using bootstrapping, in each iteration we picked two random subsets of trials, extracting for each trial the z-scored fluorescence traces between the cue onset and end of the delay period. We averaged traces across trials of each set, and calculated the pairwise Pearson correlation between the resulting two average fluorescence time-traces. We repeated this procedure 100 times, picking a different sets of random trials each time. We used one-third of the total number of *False Alarm* trials (less-frequent evaluated trial outcome) to enable the comparison with the *Hit* and *Correct Rejection* trials that occurred at higher frequency, as expected given the high session behavioral performance of the expert mice (> 80% correct trials). Miss trials were excluded from this analysis due to their infrequency (< 3%). Bouton responses were classified as stable if the correlation in response across conditions was within 90% of the inter-trial variability (correlation) across *Hits* trials used as reference.

### PCA visualization

Extracted ROI time courses were assembled in a data cube (N by S by T) of trial averaged dF/F0 responses, where N stands for the total number of cortical feedback boutons included, S is the total number of stimuli (reward-contingency trials – H, CR, FA) and T is the total number of time-bins. To reduce the dimensions of the neuronal population, this data cube was re-shaped into a data matrix (N by ST) and normalized (z-scored) such that each stimulus as a function of time represents a point in an N dimensional neural state space. To find a set of orthogonal directions that maximizes the variance captured from the data, we performed principal component analysis (PCA) and identified the *eigen* vectors of the associated covariance matrix. PCA was performed using built-in ‘*princomp*’ function in MATLAB. Data projected onto the first three principal components (PCs) is plotted in **Fig. 3c**. The variance explained by each PC is given by the ratio of its *eigen* value to the sum of all the *eigen* values. Miss trials were excluded from this analysis due to their low frequency (<3 %). On average, across fields of view, the first three PCs explained 66.1 ± 3.3 % of the signal variance for odor trials and 67.1 ± 2.8 % of signal variance explained for the sound trials.

### Machine learning based classifying task parameters and animal behavior

We used multi-layer perceptrons (MLP)^154^ to predict various stimulus and behavioral features associated with the task across trials. We normalized the datasets and further sliced them into non-overlapping windows of 4 time samples. These windows were then reshaped into a 4 * number of ROI input vectors which were given as input to the MLP. The data was split into disjoint *test* (33.3%) and *training* (66.6%) subsets respectively. For each classification problem (*stimulus*, *reward-contingency*, *instruction* and *behavior outcome*), we ran 10 repetitions with shuffling and re-splitting into *training* and *test* sets to control for variability in trial quality.

The MLP consisted of one input layer that was the same size as the input vectors, two hidden layers of 10,000 and 1,000 units each, and one output layer, whose size depended on the feature that we aimed to classify. The weights of the network were initialized using the Xavier uniform distribution^155^ (**Eq. 1**) where the *weights* of a layer (*W_ij_*) are selected from a *uniform distribution* (*U*) centered on 0 and with a range dependent on *the number of nodes* in the previous layer (*n*).

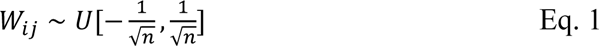

Each layer had a *Soft++* (k=1, c=2, **Eq. 2**) activation function, except for the last layer where *Softmax* normalization (**Eq. 3**) was used to return a probability distribution over the number of classes (N). For the two hidden layers we also applied a dropout rate of 50% in order to limit overfitting.

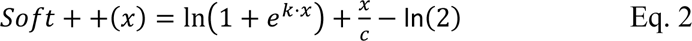

Softmax (z-*vector to be normalized*, N-*number of classes*):

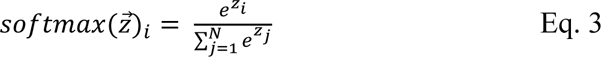

The MLP was optimized using the Adam optimization algorithm^156^, and we used as loss function the cross-entropy (**Eq. 4**) between the *Softmax* output (p) and the one-hot encoding of the label (y). Cross entropy loss (*N*-*number of classes*, y-*binary indicator* (0 or 1), if *class label c* is correct classification for observation *o*, *p*-predicted probability that observation o is of class c):

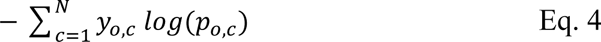

The MLP was trained for 50 epochs (iterations through the dataset) with a batch size (number of samples per step) of 25, and a fixed learning rate (weight modification rate) of 0.001, as opposed to variable learning rate which changes across epochs according to a schedule.

We ran three control analyses: *shuffle label*, *shuffle channel*, and *EGFP control*. For the shuffle label control, we shuffled the trials indices (labels) which resulted in destroying correlations between the data and the associated reward contingency. For the shuffle channel control, we shuffled the ROI fluorescence values for each time point before feeding the data into the MLP. This has the effect of destroying ROI identity information, while leaving global activity in time intact. Finally, we also ran the same analyses on EGFP data, in order to measure how much of the information in the data comes from movement related artefacts. Because EGFP fluorescence should not vary as a function of neuronal activity, any measured changes in green fluorescence are likely due to motion-related artifacts, blood-vessel occlusion or intrinsic signal changes, etc.

### Machine learning-based stimulus classification within and across blocks

Using the same procedure outlined above, we also ran a series of tests to measure how the representations of stimulus change across blocks in the experiment. For the same-block analysis, we selected only the trials from one type of block at a time (e.g., sound-go blocks) and split them in the same proportions mentioned above. For the across block analyses, we used all trials, but instead of random splitting, we assured that all trials from one type of block were in the *test* set and all trials from the other block were in the *training* set.

### Trajectory-based Kohonen mapping

Self-organized maps are a useful tool for visualizing patterns in multidimensional data, as they reduce the patterns across features to a color representation. Effectively the Kohonen mapping algorithm creates a translation key (through the model space) between a fixed color space and the data space^157,158^. This algorithm works by sequentially passing through the data samples multiple times, finding the closest match in the model space, and adjusting that model, as well as models close to it in a color space, so that they more closely resemble the sample. The color space is fixed and is three-dimensional (red, green, blue), and each model is assigned a location in this color space that does not change. The distances between models are computed in the color space, while distances between samples and models are computed in the model space. The algorithm has two parameters: the *learning rate* (which controls how large the change to the models is at each step) and the *standard deviation of a three-dimensional gaussian kernel* (which changes how many models are modified at each step, as well as how large the modification applied to those models is). We changed these parameters at each step such that at the beginning of the algorithm, many models are altered at each step (large neighborhood radius, R, **Eq. 5**) with a large learning rate (L, **Eq. 6**). This allows the algorithm to quickly make a rough estimate of the data space. Through further iterations, the radius shrinks, and the changes made are smaller, allowing the algorithm to fine-tune the models. We seeded the algorithm by selecting the 10% of samples with the lowest total activation and setting them to black. This biases the algorithm such that samples with lower activations have darker colors, and higher activations are associated with brighter colors.

The change rate and the radius of the neighborhood were given by two monotonically decreasing functions, *L(k)* and *R(k)*, respectively:

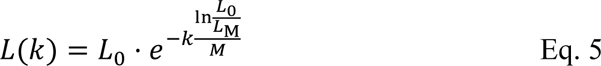

where *L(k)* is the learning rate, modulating the degree to which model vectors were changed at each training step, k. L0 and LM are initial and final learning rates. We used L0 = 1 and LM = 0.01. The total number of training steps is denoted by M.

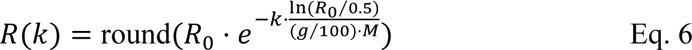

where round denotes the rounding to the nearest integer, *R(k)* specifies the neighborhood size. R0 is the initial radius of the neighborhood. g is the percentage of M after which R becomes 0. We used R0 = N/2 and g = 66 (66% of steps were used to establish the topology of the map and the remaining 34% of the steps to fine tune the representation of activity vectors in the map, i.e., only the best matching unit, BMU was changed). Within the above-defined neighborhood, model vectors move further away from the BMU changed less than the ones closer to it by multiplying the learning rate with a 3D Gaussian envelope with a SD of R(k)/3 (**Eq. 7**):

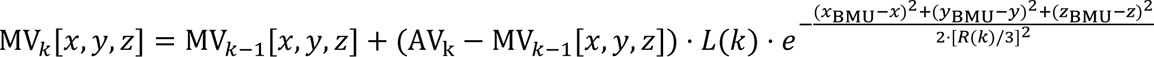

where *MVk[x,y,z]* is a model vector, at step k of the training, located within the neighborhood of the BMU [distance from BMU ≤ R(k)] at position (x,y,z) in the 3D lattice. (*xBMU*, *yBMU*, *zBMU*) denotes the position of the BMU in the 3D lattice. *AVk* is the activity vector that is learned at step k, *L(k)* and *R(k)* are the learning rate and the size of the neighborhood at step k.

### Neuronal distance across blocks

We calculated a neural distance (*1-Pearson correlation*) between the bouton ensemble response trajectory of the preceding block (averaged over responses to the last five trials in that block) and the ensemble trajectory of each trial of the current block to the same stimulus for both odor and sound trials. This metric increased systematically across trials, and matched the increase in behavioral performance in the new block *post rule-reversal* (**Figs. 4c**-**e**). Qualitatively similar results were obtained when performing the analysis for the odor and sound trials independently (data not shown).

### Optogenetic suppression of the cortical bulbar feedback

Mice were bilaterally injected at the same aPCx coordinates as for GCaMP expression with AAV to express the inhibitory opsin Jaws. On the same day, mice were bilaterally implanted with cannulas loaded with 200 μm diameter optic fibers (Doric: MFC_200/230-0.48_2.0mm_ZF1.25(G)_FLT) on top of the olfactory bulb (coordinates: AP: +1.2 from inferior cerebral vein; ML: ± 1.2 mm from inter-frontal suture; DV: 0.0 mm OB surface). The space between the optic fiber and the edges of the skull craniotomy was filled with white petrolatum (Dynarex) and the optic fiber cannulas and a metallic head-bar were attached to the skull using a combination of dental cement (Metabond®), black dental acrylic resin (Ortho-Jet™) and cyanoacrylate glue (Krazy-Glue®). After training and reaching > 80% performance, mice were ready for optogenetic suppression experiments (0.25 probability of experiencing a trial with light stimulation). Optic fiber cannulas were connected to a branched dual patch cable (Doric, Cat # SBP(2)_200/230/900-0.48_1m_SMA-2xZF1.25) using ceramic sleeves (ThorLabs, Cat # ADAL1). Light stimulation was performed using a 590 nm LED (ThorLabs Cat # M590F3) calibrated to deliver 2.4 mW at the tip of the patch cable. Stimulation (5 ms, 30 Hz light pulses) was triggered 500 ms before the start of the *cue* period and continued until the end of the *reporting* period (2.85 s). Equivalent experiments were performed in control animals expressing EGFP instead of Jaws.

