## Supplementary material for "Fast updating feedback from piriform cortex to the olfactory bulb relays multimodal reward contingency signals during rule-reversal": Hernandez_Trejo_Supplemental Information_rule_reversal

#### Supplementary Figure 1

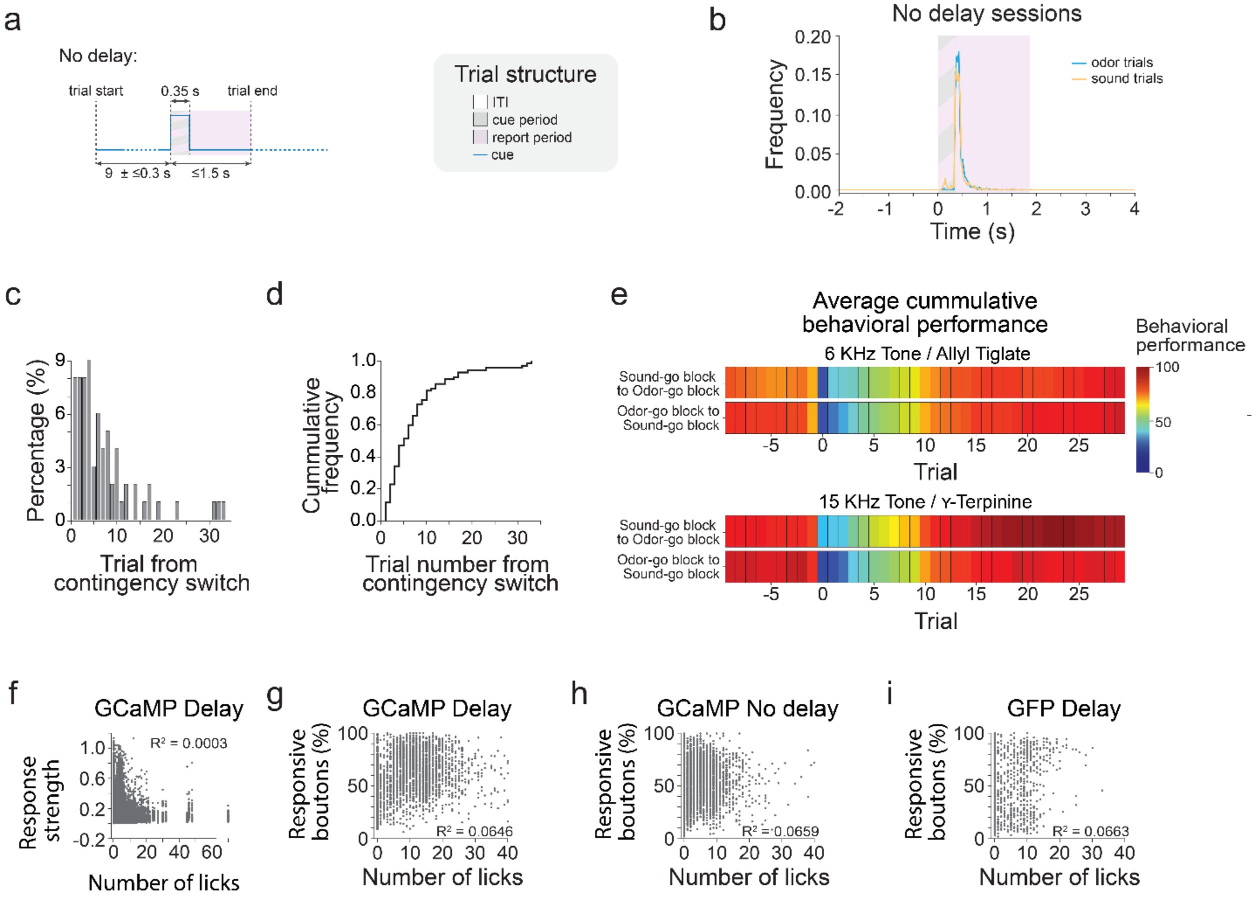


**Supplementary Figure 1**. (**a**) In the ‘no-delay’ task, a variable baseline period of (~9 ± ≤0.3 s) was followed by the delivery of a brief odor or sound cue (0.35 s) before the time when the reward became available. Mice were trained to report their decision (lick vs. no-lick) within a 1.5 s window from the cue offset. (**b**) Distributions of latency to the first-lick from cue onset from all no-delay sessions parsed by cue (blue: odor; orange: sound trials). (**c**) Quantifying switching: frequency distribution for observing five consecutive correct trials starting at the switch (rule-reversal) trial (using a 5-trials sliding window). (**d**) Cumulative distribution of the histogram from **c**. (**e**) Average behavioral performance for block transitions from example expert mice trained with multiple sound and odor cue pairs. (**f**) Number of licks versus response strength (z-score$_{max}$ – z-score$_{min}$) for GCaMP delay behavior sessions. $R^2$ = 0.0003. (**g**) Number of licks versus percentage of responsive boutons for GCaMP delay behavior sessions. $R^2$ = 0.0646. (**h**) Number of licks versus percentage of responsive boutons for GCaMP no-delay sessions. $R^2 = 0.0659$. (**f**) Number of licks versus percentage of responsive boutons for EGFP session. $R^2 = 0.0663$.

#### Supplementary Figure 2

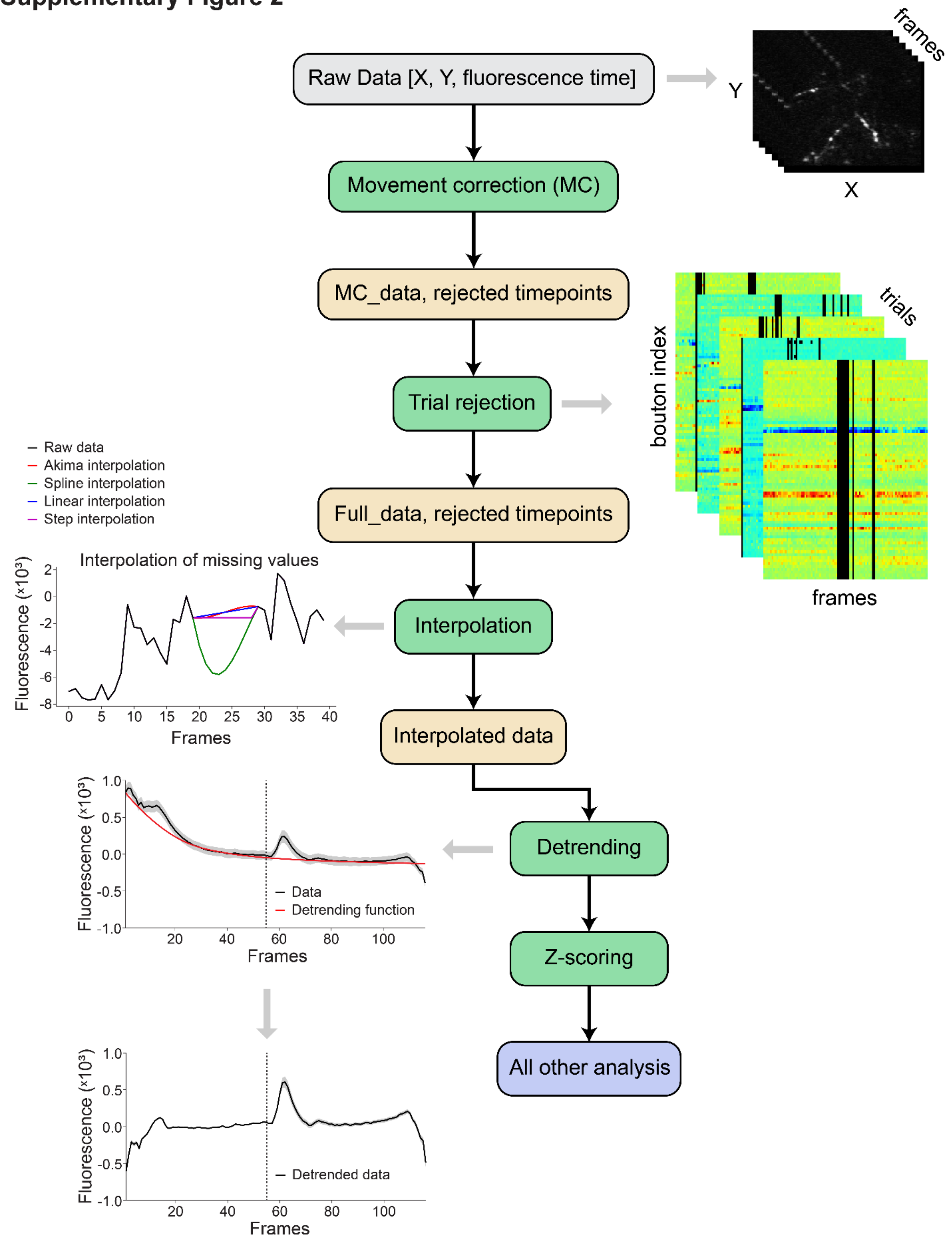

**Supplementary Figure 2**. Outline of the data pre-processing pipeline which includes the following steps: movement correction, region of interest (ROI) selection, trial rejection, interpolation, de-trending and z-scoring (Methods).

### Supplementary Figure 3

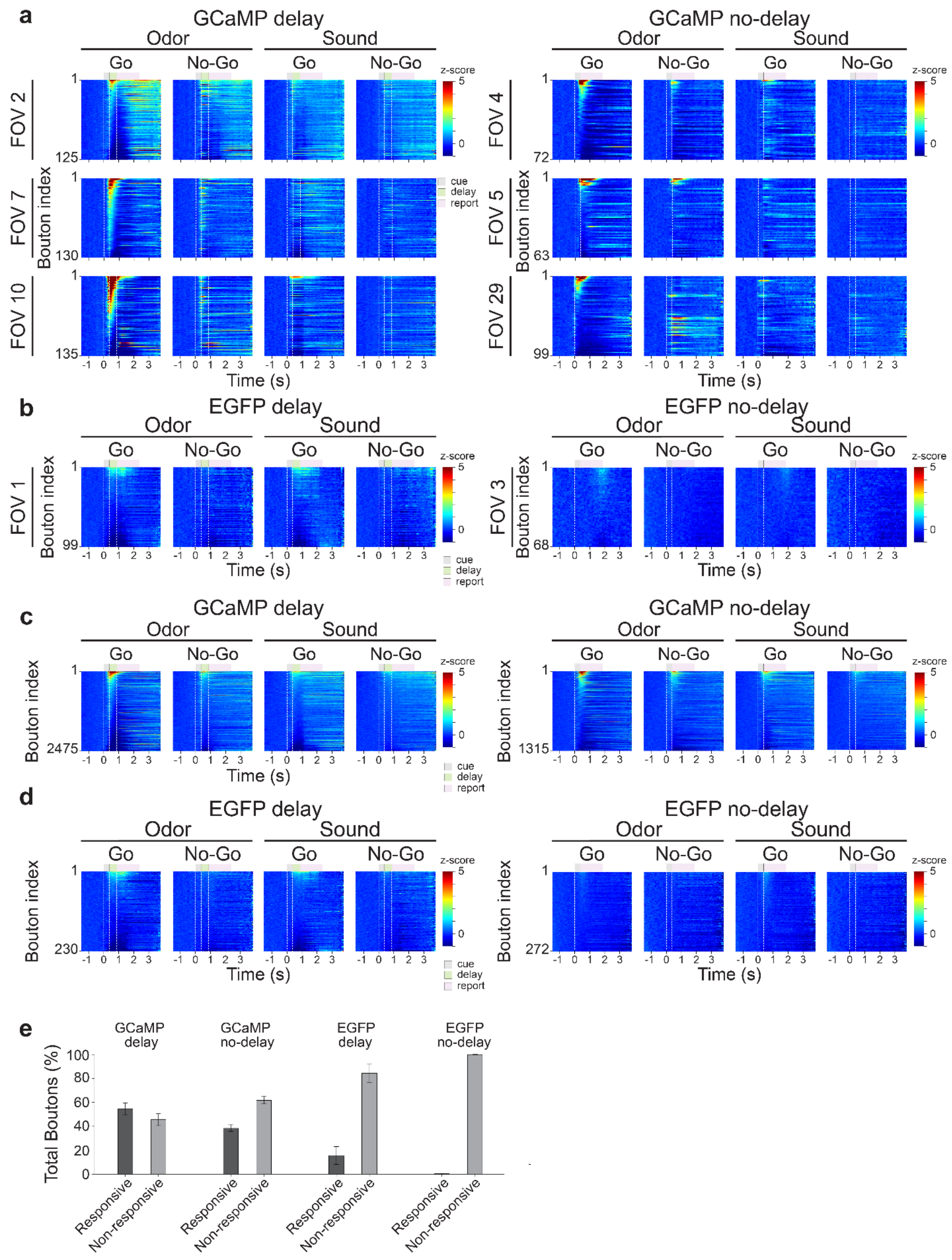

**Supplementary Figure 3**. (**a**) Example trial-averaged GCaMP5 bouton responses from *delay* (*Left*) and *no-delay* (*Right*) behavior and imaging sessions in expert mice sorted by the cue identity (Odor and Sound) and instruction signals (Go and No-Go). (**b**) Same for example trial-averaged EGFP bouton fluctuations from *delay* (*Left*) and *no-delay* (*Right*) sessions. (**c**) Same as **a** for all boutons (GCaMP5) across all fields of view in the *delay (Left)* and *no-delay* (*Right*) versions of the task. (**d**) Same as **c** for the EGFP boutons. (**e**) Percentage of boutons classified as 'Responsive' or 'Non-responsive', from all GCaMP and EGFP imaging and behavior sessions in expert mice trained in both versions of the task (Methods).

#### Supplementary Figure 4

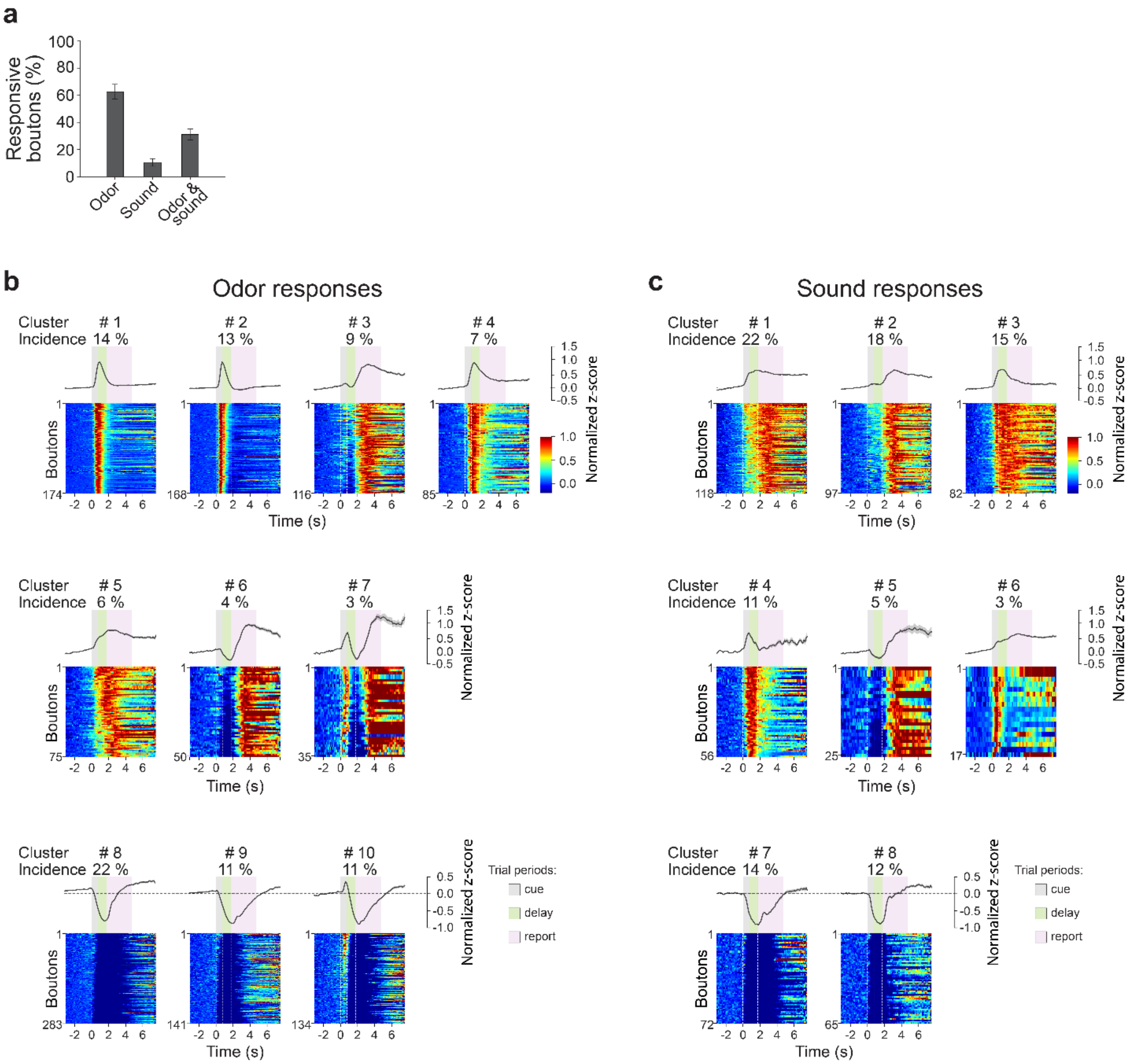


**Supplementary Figure 4**. (**a**) Percentage of responsive boutons to the odor, sound, or both cues across all fields of view in the delay version. (**b**,**c**) K-means clustering of bouton responses for odor trials (**b**) and sound trials (**c**) in the *delay* version of the task. Average cluster waveform (*enhanced*, *suppressed*, *complex*) and cluster incidence (*Top*) and individual bouton response (*Bottom*) for each cluster. Note the diversity of time courses across clusters, spanning different periods of the task. Trial periods (*cue*, *delay*, *report*) are marked by vertical lines and colored areas.

#### Supplementary Figure 5

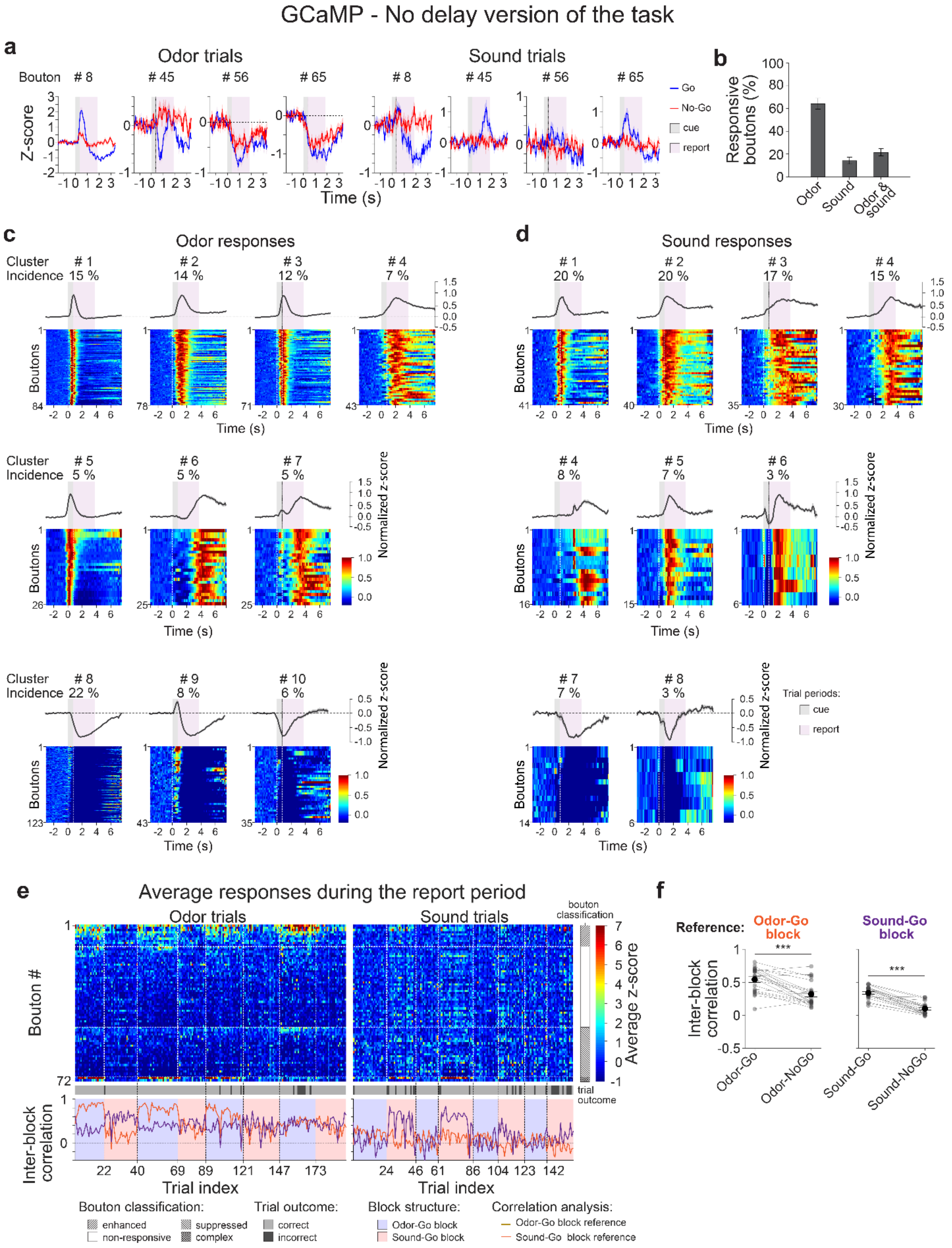

**Supplementary Figure 5**. *GCaMP no-delay sessions*: (**a**) Example average bouton responses to odor (*Left*) and sound (*Right*) cues sorted by instruction signals (Go, *blue* and No-Go, *red*). Trial periods are marked by colored areas (light gray: *cue*; light pink: *report*). (**b**) Percentage of responsive boutons to the odor, sound, or both cues. (**c**,**d**) K-means clustering of GCaMP responses in *no-delay* sessions for odor (**c**) and sound trials (**d**). Average cluster waveform (*enhanced*, *suppressed*, *complex*) and cluster incidence (*Top*) and individual bouton response (*Bottom*) for each cluster. Trial periods are marked by vertical lines and colored areas. (**e**) Activity maps (*Top*): Average responses (z-scored) during the *report period* for all identified boutons from an example behavior and imaging session across trials. Odor trials (*Left*) are parsed from sound trials (*Right*). *Bouton classification*: Bouton responses are grouped depending on their responsiveness and polarity and classified as enhanced (*right oblique pattern*), suppressed (*left oblique pattern*), or complex (*cross pattern*). A subset of boutons was classified as unresponsive (*no pattern*). Color scale bar: Average z-score values. Bottom bar represents the outcome of each trial. Correct trials (*hits* and *correct rejections*) are shown in *light gray*. Incorrect trials (*misses* and *false alarms*) shown in *dark gray*. Bottom plots: Average correlation coefficient between an average bouton ensemble response vector (Methods) of the first 'Odor-Go block' (orange trace) or first 'Sound-Go block' (purple trace) in the session, and the ensemble bouton response vector of each trial parsed by cue identity. Block structure: 'Odor-Go blocks': *light purple*; 'Sound-Go blocks': *pink* parsed by cue. (**f**) Average inter-block correlation coefficients for all fields of view (*no-delay* version) using the 'Odor-Go block' (*Left*) or 'Sound-Go block' (*Right*) as reference for the ensemble bouton responses during the odor and sound trials and for both types of instructions. Student's t-test: *** = $p < 0.0001$; n.s.: non-significant. All panels error bars: ±SEM.

#### Supplementary Figure 6

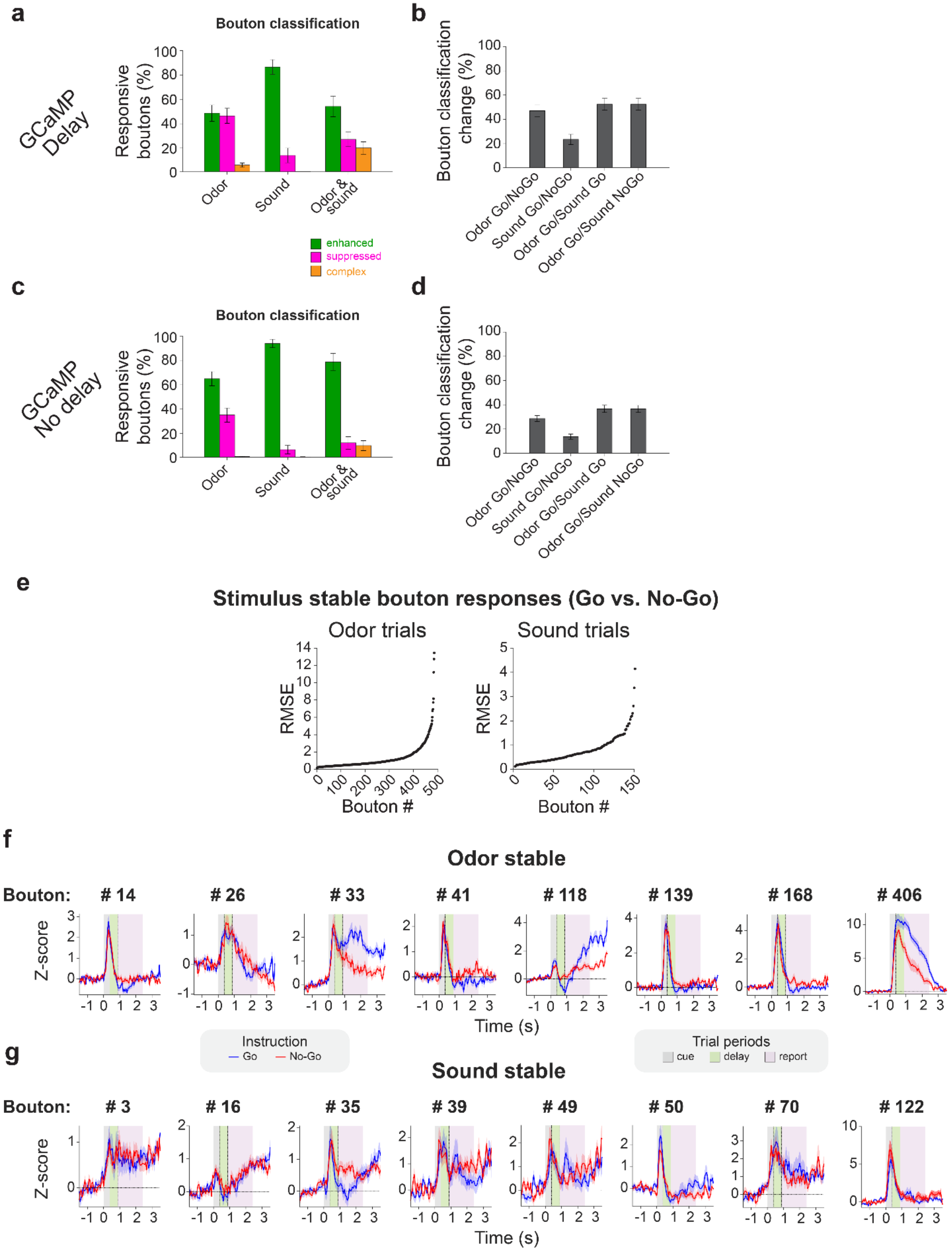

**Supplementary Figure 6:** (**a**,**c**) Percentage of GCaMP labeled boutons classified as enhanced (*green*), suppressed (*magenta*) or complex (*orange*) for the *delay* (**a**) and *no-delay* (**c**) versions of the task. (**b**,**d**) Percentage of boutons that changed class (e.g. from unresponsive to enhanced, enhanced to suppressed, etc.) in *delay* (**b**) and *no-delay* (**d**) when comparing responses to the same sensory cue for different instructions (Go vs. No-Go) and to different cues (odor or sound) for the same instruction. (**e**) Root mean square errors (RMSE) between average responses of individual boutons (during the *cue* and *delay* periods) in Go versus No-Go trials for same sensory stimulus; shown are RMSE for boutons that did not substantially change their response waveform *across* conditions (i.e. Pearson correlation coefficient between individual bouton response waveforms in the Go vs. No-Go trials for the same sensory cue was within the 90% percentile of the distribution of trial-to-trial correlations for the Go trials). RMSE were sorted by increasing magnitude for odor responsive (*Left*) and sound responsive (*Right*) boutons that passed the criterion. (**f**,**g**) Example odor (**f**) and sound (**g**) responses of boutons across signal instructions (Go vs. No-Go) from the distributions shown in (**e**). Numbers (#) represent indices of the corresponding boutons in the RMSE distributions.

#### Supplementary Figure 7

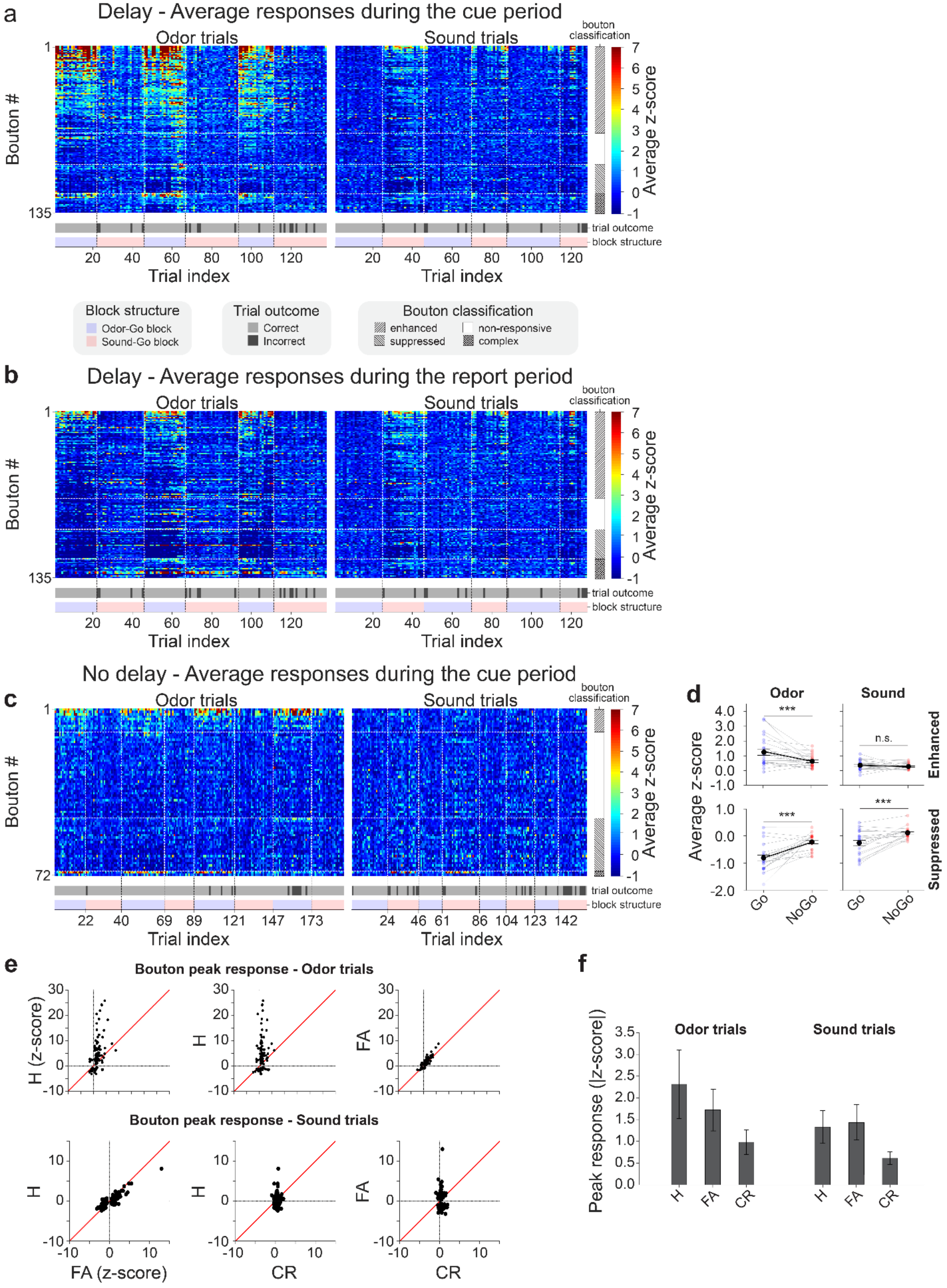

**Supplementary Figure 7**. (**a**,**b**) Activity maps for the same field of view shown in **Figs. 2d,e** representing average responses (z-scored) during *cue* (**a**) and *report* periods (**b**). (**c**) Activity map along trials of average responses (z-scored) during the *cue* period in a *no-delay* example session. (**d**) Average z-scored response values during the *cue* period (*no-delay* sessions) parsed by cue (Odor or Sound) and instruction (Go or No-Go) for the responsive boutons sampled. Each pair of connected dots represents average z-scored response values computed across boutons from one behavior session. Black dots represent average z-scored ensemble bouton response across sessions. (**e**) Peak responses (z-scored) across different behavioral contingencies (Hits vs. False alarms, Hits vs. Correct rejections and False alarms vs. Correct rejections) of all responsive boutons in an example field of view. Each dot corresponds to a bouton. (**f**) Average peak responses (z-scored) across different behavioral contingencies (Hits, H, False alarms, FA, Correct rejections, CR) across fields of view (*delay* sessions) for Odor (*Left*) and Sound (*Right*) trials. Student's t-test: *** = $p < 0.0001$; n.s.: non-significant. All panels error bars: ± SEM.

### Supplementary Figure 8

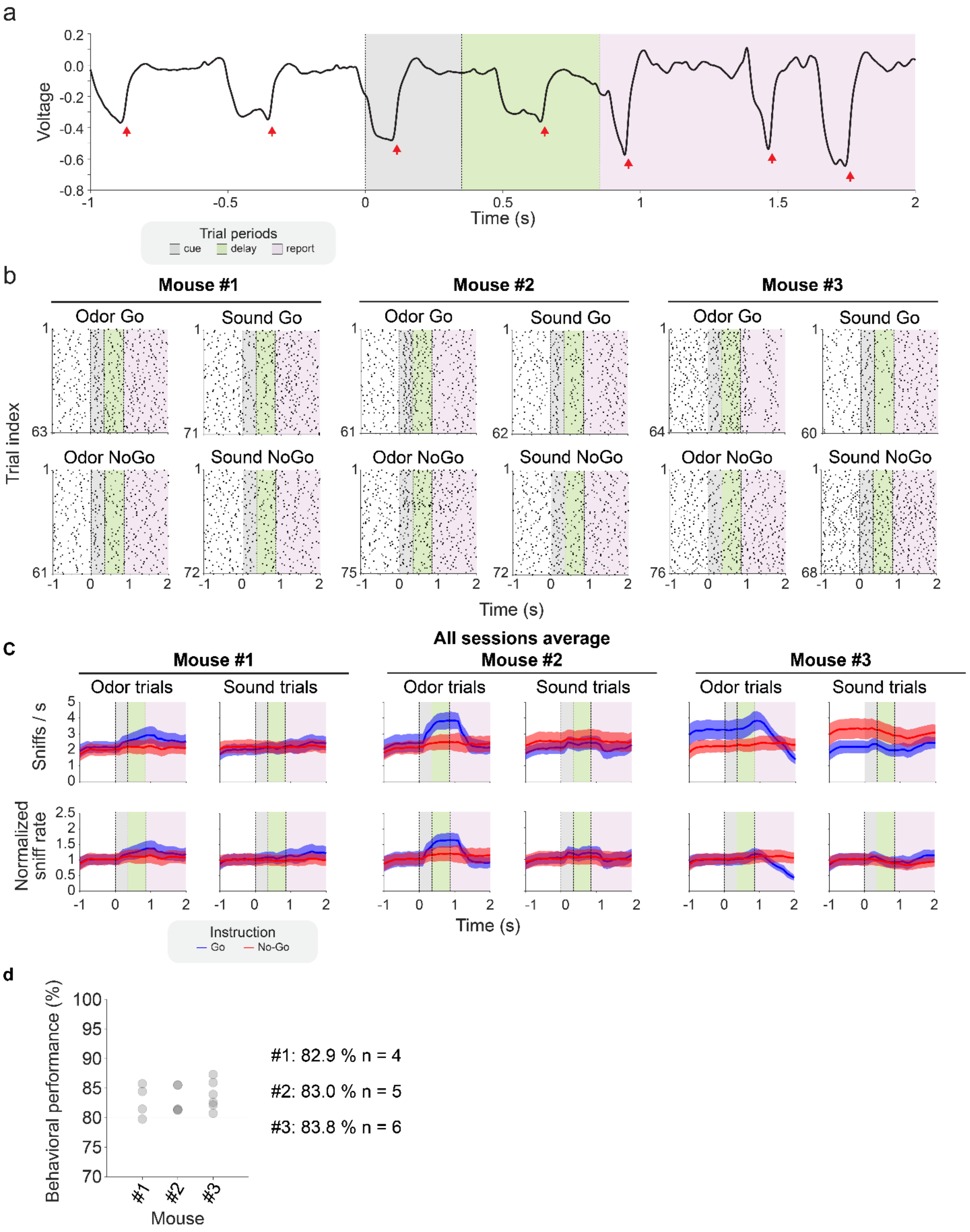

**Supplementary Figure 8**. Sniff patterns in control expert mice performing the *delay* version of the rule-reversal task. (**a**) Example sniff pattern during one trial in an expert mouse. Sniffing was measured using a flow sensor placed in front of the animal's snout (Methods). Arrows correspond to inhalation start. (**b**,**c**) Sniff monitoring across trials in example behavior sessions. For each mouse, trials are sorted by cue identity (Odor vs. Sound) and instruction signal (Go vs. No-Go). (**b**) Each dot represents an inhalation start identified as shown in **a** (Methods). Colored areas represent different trial periods (*cue*, *delay*, *report*). (**c**) Average sniff rate (*Top*) and normalized average sniff rate (*Bottom*, with respect to pre-cue air baseline) across all sampled behavior sessions (4-6 sessions per mouse). Shaded area - SEM (**d**) Average behavior performance of the mice shown in (**b**,**c**) (Mouse 1 - 82.9%, n= 4 sessions; Mouse 2 - 83.0%, n= 5 sessions; Mouse 3 - 83.8%, n= 6 sessions). Across mice and sessions, sniff rate for a given cue was either unaffected or changed slightly based on the instruction signal. Instruction-dependent changes in sniff rate varied across animals and did not appear to correlate with the behavioral performance.

#### Supplementary Figure 9:

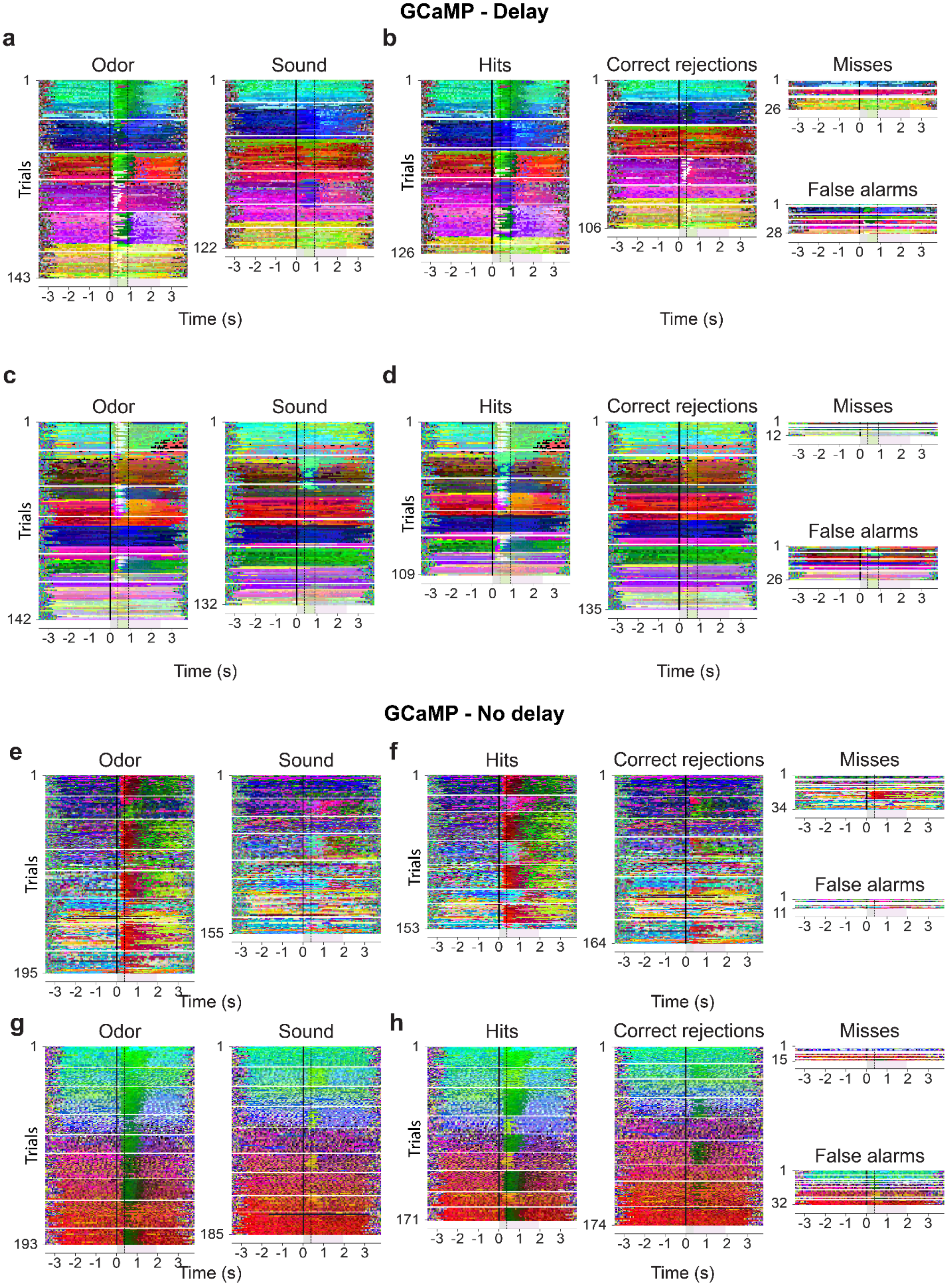

**Supplementary Figure 9**. Kohonen response maps of example *delay* (**a-d**) sessions. (**a,c**) Example fields of view #7 (**a**) and #10 (**c**) parsed by *cue* (Odor vs. Sound). (**b**,**d**) Example fields of view #7 (**b**) and #10 (**d**) parsed by trial contingency (H, FA, CR). (**g-h**) Same analysis for example fields of view in the *no-delay* version of the task.

#### Supplementary Figure 10

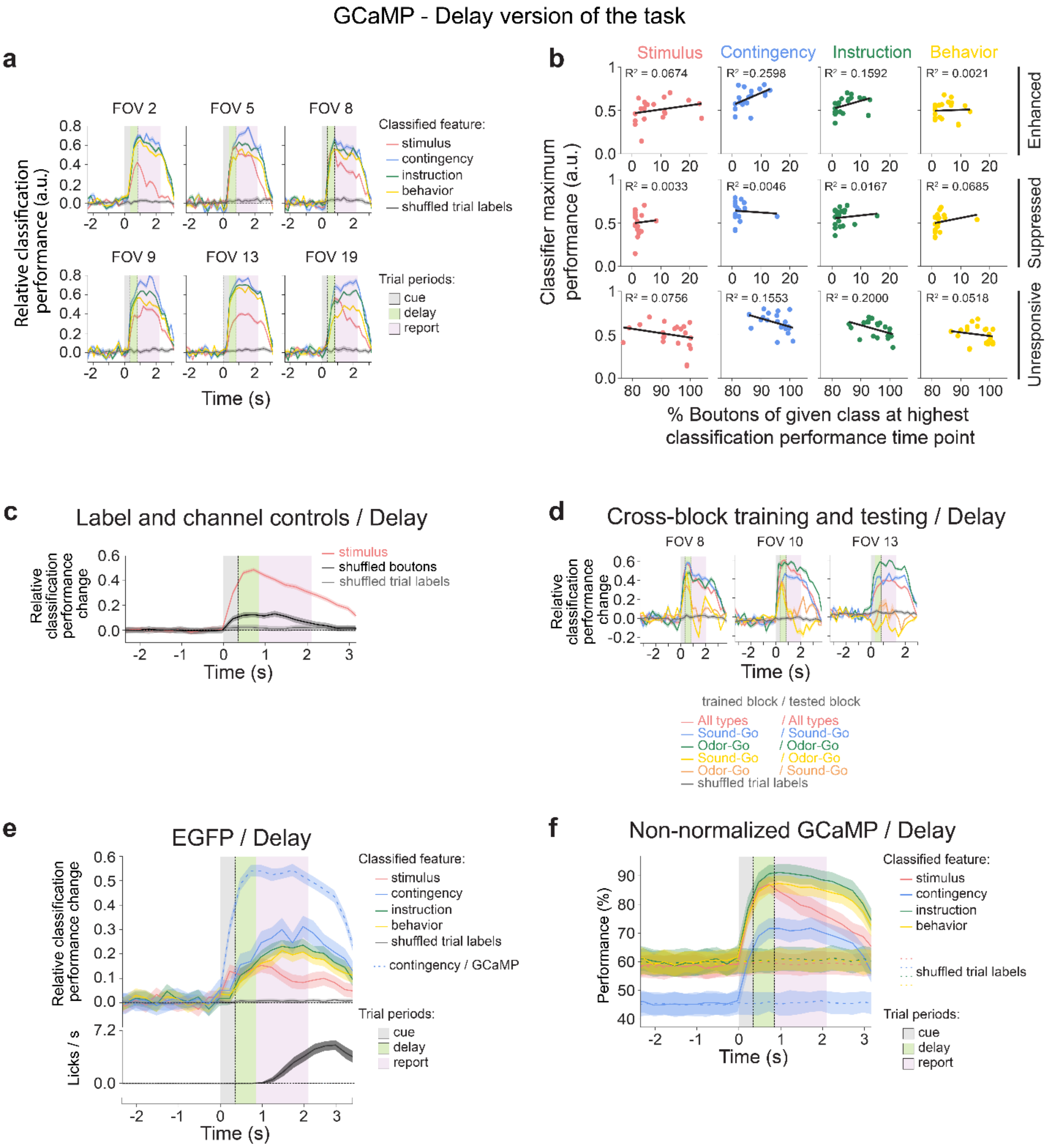


**Supplementary Figure 10**. GCaMP *delay* sessions: (**a**) Performance of multilayer-perceptron classifiers (MLP, **Fig. 3d**) for example individual fields of view. MLP were trained to decode stimulus identity (odor or sound), behavioral contingency (H, FA, CR), trial instruction (Go or No-Go), and behavior (lick or no-lick). When shuffling trial labels on the training data, the average classifier performance was 0. (**b**) Pearson correlation analysis between the percentages of boutons classified as enhanced (*Top*), suppressed (*Middle*), or unresponsive (*Bottom*) and the maximum performance of the multilayer-perceptrons (**Fig. 3d**). (**c**) Multilayer-perceptron classifier performance for stimulus classification substantially decreases when shuffling the bouton indexes or the trial labels of each trial. (**d**) Cross-block training and testing: Multilayer-perceptron performance for classifying *cue* identity (Odor vs. Sound) when training using Odor-Go and Sound-Go block trials, Odor-Go block trials only, or Sound-Go block trials only. Performance is tested either on *same* type of block as training or *across* block types (e.g. train on Odor-Go block trials and test on Odor-Go block trials; train on Odor-Go blocks trials only and test on Sound-Go trials, etc.) (**e**) Multilayer-perceptron performance for classifying stimulus, contingency, instruction and behavioral outcome, using boutons expressing EGFP in the cortical-bulbar feedback instead of GCaMP. GCaMP classifier performance at predicting trial contingency (H, FA, CR) is shown as reference (light blue line). When shuffling trial labels on the training data, average classifier performance was 0. (*Bottom*) Distribution of number of licks per second across all sessions. (**f**) Un-normalized version of the multilayer-perceptron performance for classifying stimulus, contingency, instruction and behavioral outcome (GCaMP – delay version of the task) for comparison to the normalized version shown in **Fig. 3d**.

#### Supplementary Figure 11

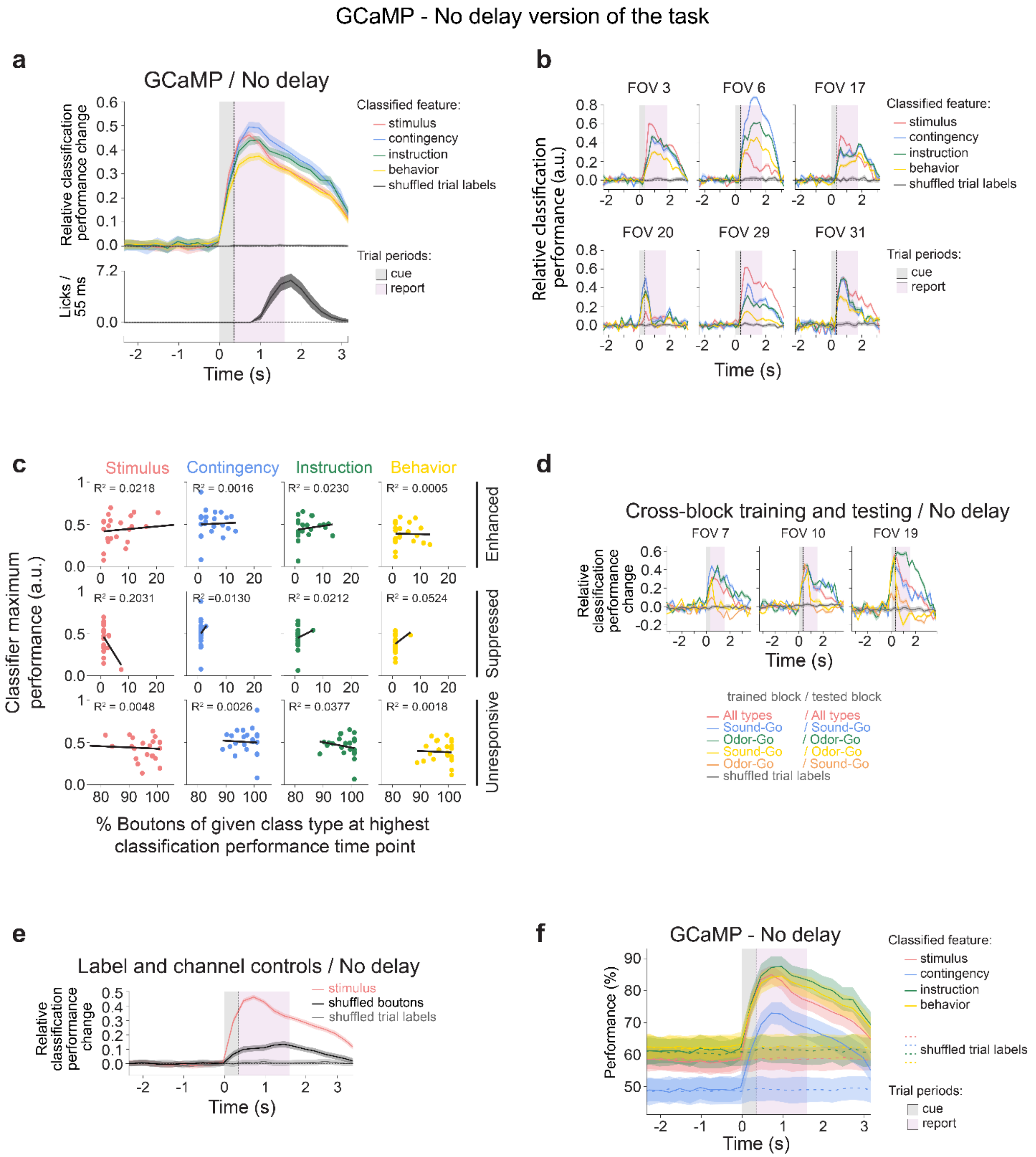


**Supplementary Figure 11.** GCaMP *no-delay* sessions. (**a**) Multi-layer perceptrons were trained to decode stimulus identity (Odor or Sound), behavioral contingency (Hit, FA, CR, or Miss), trial instruction (Go or No-Go), and behavior (lick or no-lick). *Top:* Cross-sessions average classifier performance normalized relative to the baseline. Shuffling the labels on the training data results in baseline performance of the classifier. (*Bottom*) Frequency distributions of licks per second for all acquired sessions. (**b**) Multilayer-perceptron performance for example fields of view. (**c**) Pearson correlation analysis between the percentage of boutons classified in a field of view as enhanced (*Top*), suppressed (*Middle*), or unresponsive (*Bottom*) and the maximum performance of the multilayer-perceptrons from **a**. (**d**) Cross-block training and testing: Multilayer-perceptron performance for classifying *cue* identity (Odor vs. Sound) when training using Odor-Go and Sound-Go block trials, Odor-Go block trials only, or Sound-Go block trials only. Performance is tested either on *same* type of block as training, or *across* block types (e.g. train on Odor-Go block trials and test on Odor-Go block trials; train on Odor-Go blocks trials only and test on Sound-Go trials, etc.) (**e**) Multilayer-perceptron performance for stimulus classification when shuffling the bouton channel indexes and the labels of each trial. (**f**) An un-normalized version of the multilayer-perceptron performance shown in **a**.

### Supplementary Figure 12

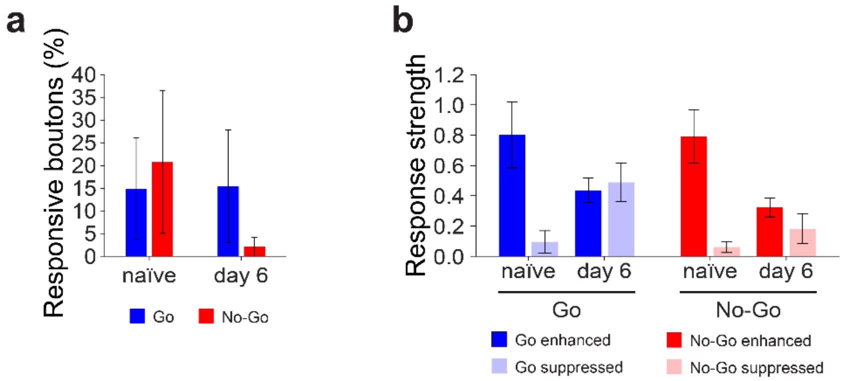


**Supplementary Figure 12**. Two-Sounds Go/No-Go task: (**a**) Percentage of responsive boutons in Go and No-Go trials in naïve sessions and after six days of training. (**b**) Average bouton response strength ($dF/F_0$) in naïve mice and after six days of training. All panels error bars: ± SEM.
